# Glioblastoma Tumors with Decelerated Epigenetic Aging Are Characterized by Glutamatergic Neuronal Activity and Stemness

**DOI:** 10.64898/2026.08.29.747960

**Authors:** Meysam Motevasseli, Majid Eterafi, Hananeh Alaei, Pouyan Zandi, Neda Shajari, Mina Tabrizi, Elham Safarzadeh

## Abstract

**Introduction:** Gliomas integrate into neural circuits and heighten neuronal excitability, engaging in bidirectional communication whereby neuronal activity promotes tumor growth and proliferation. Aging reshapes the brain microenvironment through extracellular matrix changes, altered secretory factors, and immune dysfunction, creating conditions permissive to tumorigenesis and limiting immunotherapy efficacy in glioblastoma. However, its effect on neuronal excitability and signaling in glioblastoma remains poorly understood.

**Methods:** We developed a novel classification system for glioblastoma by leveraging three classes of DNA methylation-based aging biomarkers: chronological, biological, and mitotic clocks. This approach stratified tumors into accelerated and decelerated epigenetic aging subtypes, which we then characterized at the molecular, functional, and clinical levels using multimodal analyses. Guided by these profiles, we evaluated the in vitro effects of the FDA-approved agents levetiracetam and riluzole, alone and in combination with temozolomide, on U87MG and A172 cell lines. Specifically, we assessed changes in cell viability, apoptosis, and the expression of marker genes related to stemness, neuronal hyperexcitability, and immunosuppression.

**Results:** Tumors with decelerated epigenetic aging showed expression modules and CpG hypomethylation associated with neuronal activity and stemness, and carried significantly worse prognosis. Single-cell and spatial multi-omics analyses revealed enrichment for neurons and malignant neural stem-like cells in these tumors. They also displayed enhanced intercellular communication, driven predominantly by glutamate signaling across the malignant, neuronal, and immune compartments of the tumor microenvironment. In vitro pharmacological inhibition of glutamatergic signaling with levetiracetam and riluzole reduced cell viability, induced apoptosis, and suppressed expression of stemness, neuronal hyperexcitability, and immunosuppression markers. Both agents potentiated the cytotoxic and apoptotic effects of temozolomide, supporting glutamatergic inhibition as a strategy for improving chemosensitivity.

**Conclusion:** By establishing a framework for decoding glioblastoma heterogeneity through epigenetic aging, we identified the glutamatergic pathway as a clinically actionable vulnerability. Our findings suggest that combining anti-glutamatergic therapies with temozolomide exerts synergistic antitumor effects while mitigating adverse chemotherapy-induced phenotypes, such as increased stemness, neuronal hyperexcitability, and immunosuppression, thereby laying the groundwork for novel therapeutic strategies.

**Graphical Abstract:** 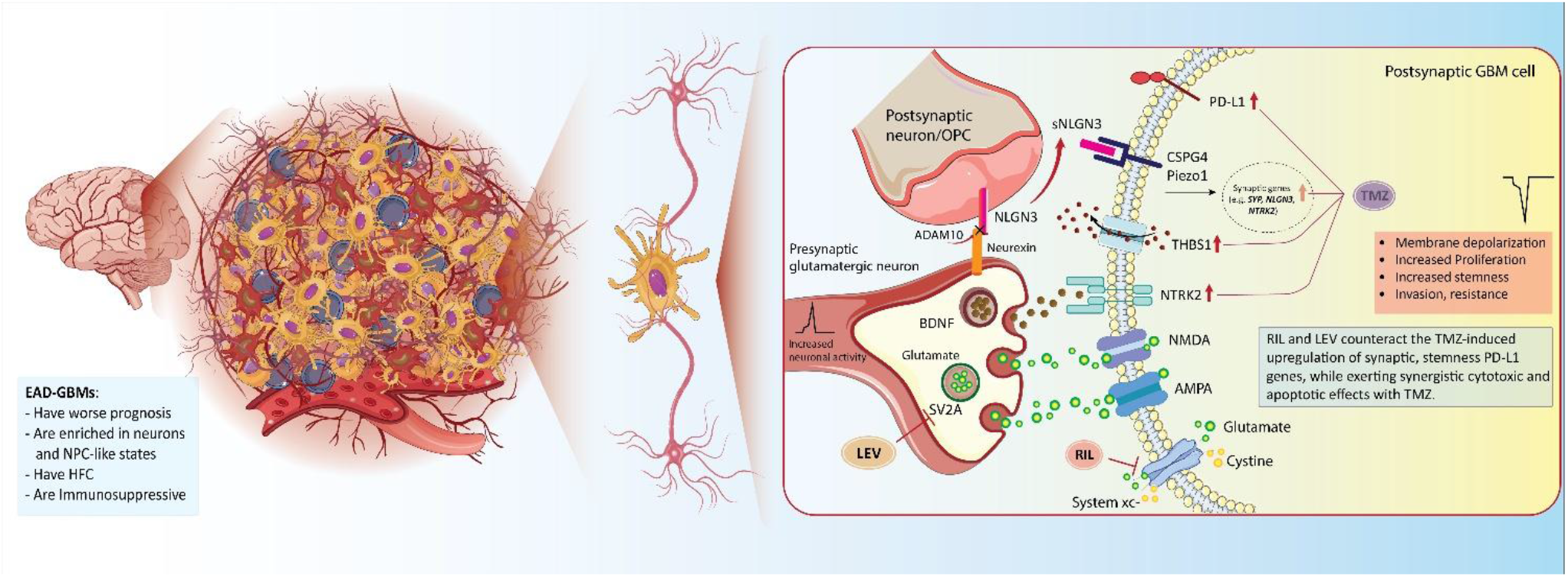

## Introduction

As the field of cancer neuroscience advances, it has become evident that neurons play a crucial role in development and progression of glioblastoma (GBM), the most common and aggressive primary brain malignancy. GBM constitutes approximately 50.9% of all brain malignancies and continues to carry a dismal prognosis, despite advances in the surgical and medical treatments [1]. Previous research has shown that neurons form bona fide glutamatergic synapses with glioma cells, mediated by α-amino-3-hydroxy-5-methyl-4-isoxazolepropionic acid (AMPA) receptors, which drive tumor progression and invasion [2]. Additionally, gliomas electrically integrate into neural circuits through both synaptic connections and gap junction-mediated networks [3]. Neuroligin-3 (NLGN3) promotes glioma growth by activating multiple oncogenic pathways and inducing its own feed-forward expression, enhancing tumor proliferation and synapse formation [4]. Brain-derived neurotrophic factor (BDNF) facilitates glioma progression by strengthening neuron-to-glioma synaptic connections through TrkB (encoded by *NTRK2*) signaling and AMPA receptor trafficking, both representing key neuronal activity-dependent paracrine signaling factors [5]. Recent findings highlight thrombospondin-1 (THBS1) as a central factor promoting glioma-neuron crosstalk in intratumoral regions of GBM characterized by high functional connectivity (HFC) associated with poorer survival [6].

Although GBM develops in individuals of all ages, including children and adults, the age- and gender-adjusted incidence is highest among older adults aged 75 to 84 years [7]. While individuals over 70 represent an increasing proportion of GBM cases [8], they are often excluded from clinical trials due to their poor prognosis and limited survival (6-9 months) [9]. Older GBM patients frequently exhibit distinct genomic and epigenomic signatures compared to their younger counterparts [10]. Aging transforms the microenvironment by altering extracellular matrix composition, secretory factors, and immune system functionality. Such changes create a permissive niche for tumorigenesis [11]. Notably, neural progenitor cells (NPCs) undergo critical changes with aging [12]. Older age also enhances immunosuppression in the brain and reduces the effectiveness of immunotherapy in GBM patients [13], contributing as a limiting factor to successful CAR-T cell responses [14]. Brain aging also significantly impacts neuronal excitability and glutamate signaling through multiple interconnected mechanisms. Physiological brain aging is characterized by decreased glutamatergic transmission, primarily due to reduced N-methyl-D-aspartate (NMDA) receptor density and impaired glutamate uptake capacity [15]. Structural changes include reduced dendritic complexity and spine numbers, leading to decreased spontaneous glutamate receptor-mediated responses while paradoxically increasing action potential firing rates [16].

Growing evidence suggests that epigenetic changes contribute significantly to cancer development and progression. Age-dependent changes in the methylation of specific CpG sites can be measured and represented as epigenetic age (EA) [17]. DNA methylation (DNAm) clocks, have emerged as powerful tools for estimating biological age, which is linked to various age-related diseases, including cancer [18]. In glioma, EA is a promising factor for elucidating tumorigenesis, predicting cancer risk, and assessing patient outcomes [19, 20]. Their inclusion in clinical trials has been proposed as a strategy to optimize treatments that preserve both lifespan and health span [21]. During physiological brain aging, global DNA hypomethylation and site-specific promoter hypermethylation occur in genes regulating synaptic plasticity, glutamate signaling, and neurogenesis. These epigenetic changes contribute to neurobiological decline, which is characterized by impaired cognition, increased neuronal vulnerability, and a disturbance in the excitatory-inhibitory balance [22]. Therefore, age-dependent GBM immunosuppression and plasticity may arise from intricate interactions among neurons, immune cells, and malignant tumor cells; yet, the underlying mechanisms governing this crosstalk remain incompletely understood. This research gap is critical to address, as finding direct causal evidence can unlock new therapeutic opportunities.

In this study, we applied multiple EA markers to bulk DNAm data, to classify GBM patients into EA accelerated (EAA) and decelerated (EAD) groups, which were subsequently characterized at molecular, functional, and clinical levels using multi-omics analyses. Our findings revealed that glutamate signaling, originating mainly from neuronal and stem-like compartments of GBM, is markedly elevated in EAD cases. In vitro pharmacological inhibition of glutamatergic excitatory signaling using the FDA-approved drugs levetiracetam (LEV) and riluzole (RIL) significantly reduced the expression of genes associated with stemness, neuronal hyperexcitability, and immunosuppression. Furthermore, these drugs demonstrated synergistic cytotoxic and apoptosis-inducing effects when combined with temozolomide (TMZ).

## Results

### Epigenetic aging clocks dichotomize GBM patients

We examined age-related DNAm patterns in IDH-WT GBM patients by employing the publicly available TCGA-GBM Illumina 450K tissue DNAm dataset from The Cancer Genome Atlas (TCGA; n=126). Since most EA clocks rely on penalized linear regression, they can miss biologically important patterns driven by complex, non-linear relationships. To better capture the multifaceted biology of aging, we employed multiple classes of DNAm clocks: chronological, biological, and mitotic, and examined their distribution patterns across GBM patients (Figure. 1A, B).

**Figure 1.**
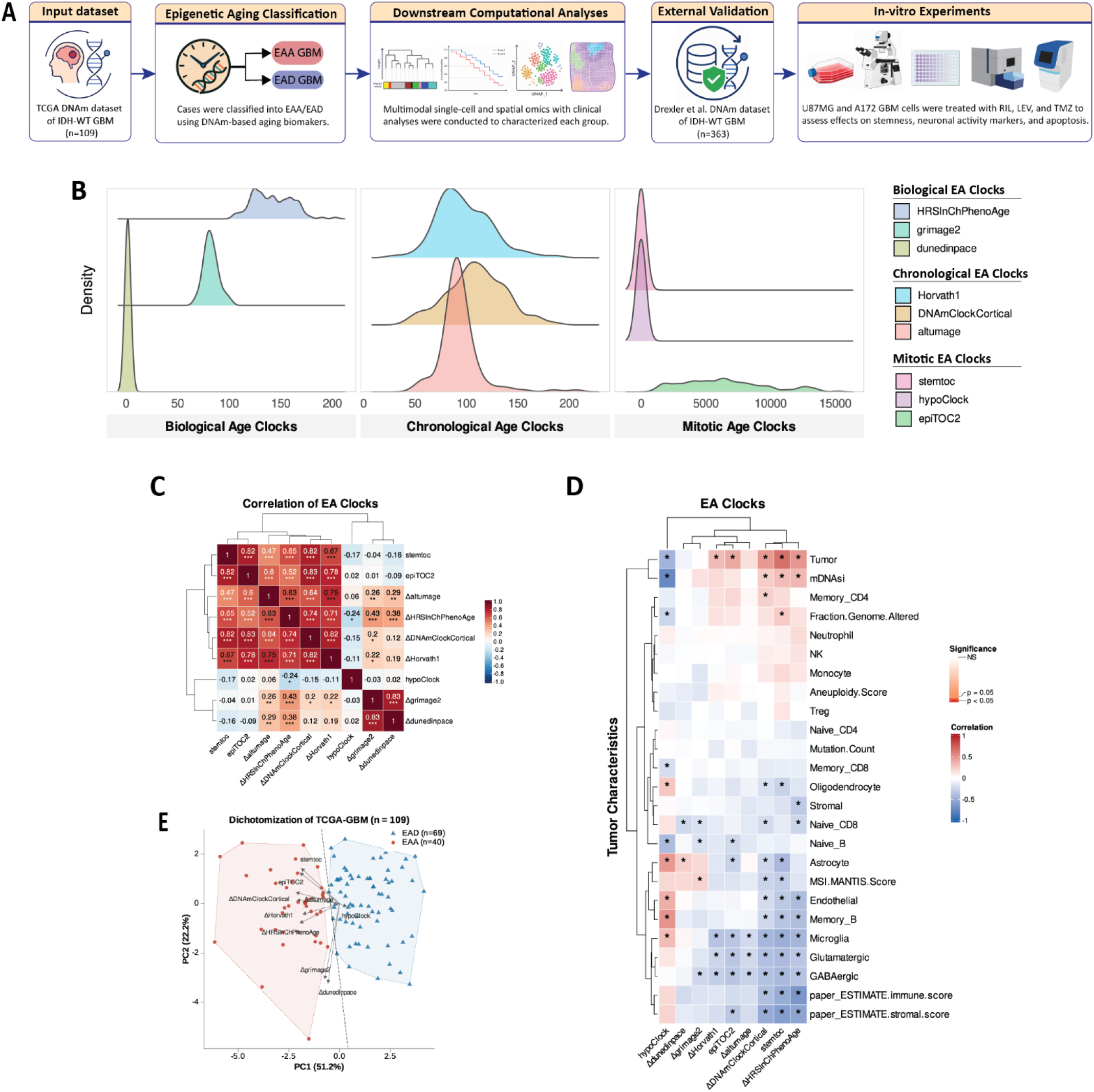
EA clocks in IDH-WT GBM: associations with tumor characteristics and patient stratification. **(A)** Schematic of the study workflow. In patients diagnosed with IDH-WT GBM (n=109), we calculated EA clocks using bulk-tissue DNAm arrays and stratified cases into EAA and EAD groups for downstream analyses. (**B**) Density plots showing the distribution of the calculated EA clocks across GBM cases. (**C**) Heatmap plot depicting correlations among EA clocks. (**D**) Heatmap plot showing correlations between EA clocks and deconvoluted tumor cell populations and characteristics. (E) PCA-LDA plot illustrating the classification of cases into EAA and EAD groups.

Chronological DNAm clocks were developed to accurately quantify chronological age (CA) in blood or multiple tissues based on patterns at specific CpG sites and assess the effects of longevity interventions. Two widely used clocks, Horvath1 and AltumAge, are trained on multiple tissues [23]. A recent brain cortex-specific chronological clock, known as the Cortical clock, optimally predicts age of the human cortex [24]. Second-generation EA markers, known as biological clocks, combine CA with composite clinical indicators, significantly improving the prediction of diverse age-related outcomes such as mortality risk, cancer onset, and Alzheimer’s disease. PhenoAge, for instance, estimates phenotypic age by combining DNAm at 513 CpG sites with nine clinical biomarkers: albumin, creatinine, glucose, C-reactive protein, lymphocyte percentage, mean corpuscular volume, red cell distribution width, alkaline phosphatase, and leukocyte count. HRSInChPhenoAge is the updated PhenoAge which expands the original 9-marker PhenoAge to include more comprehensive clinical measures of functional and cognitive ability [25]. GrimAge2 is a second-generation DNAm biomarker trained on plasma-protein surrogates and smoking-related measures that improves mortality prediction [26], while DunedinPACE is a next-generation DNA-methylation biomarker of the pace of aging derived from longitudinal organ-system decline data [27]. We also included mitotic clocks, which estimate the cumulative number of cell divisions in tissues; a process associated with cancer risk. Notable models include epiTOC2, HypoClock, and stemTOC. EpiTOC2 accurately estimates stem cell division rates in normal tissues and has been shown to outperform HypoClock in discriminating preneoplastic lesions. HypoClock relies specifically on hypomethylation patterns at solo-WCGW sites [28]. Furthermore, stemTOC, an improved pan-tissue DNAm counter, reflects increased mitotic age in precancerous lesions and normal tissues exposed to cancer risk factors [29].

Epigenetic age acceleration (EAA, denoted as Δ) was calculated for chronological and biological clocks by subtracting CA from EA. This calculation was not applied to mitotic clocks, as they measure cumulative cell divisions rather than acceleration relative to chronological age; therefore, raw values were used for these clocks in subsequent analyses. Correlation analysis of the EA clocks revealed four distinct clusters (Figure 1C). The first cluster comprised ΔGrimAge2 and ΔDunedinPACE, which exhibited a strong positive correlation (r = 0.83, p < 0.0001). HypoClock formed the second cluster in isolation. The third cluster included ΔAltumAge, ΔHRSInChPhenoAge, ΔDNAmClockCortical, and ΔHorvath1. Finally, the two mitotic clocks — stemTOC and epiTOC2 — constituted the fourth cluster, with inter-clock correlations reaching r = 0.82 (p < 0.0001).

To investigate how different EA clocks capture tumor features in GBM, we performed correlation analyses between these clocks and tumor characteristics. The analysis revealed distinct clustering patterns of both EA clocks with tumor microenvironment (TME) features. Several EA clocks, including ΔDNAmClockCortical, stemTOC, and ΔHRSInChPhenoAge, were positively correlated with tumor purity and DNAm-based stemness index (mDNAsi), while showing significant negative correlations with deconvoluted brain-resident cell populations (microglia, glutamatergic neurons, and GABAergic neurons), ESTIMATE immune and stromal scores. In contrast, hypoClock displayed an opposing pattern, with negative correlations with tumor purity and mDNAsi, and positive correlations with oligodendrocytes, astrocytes, endothelial cells, memory B cells, and microglia. In GBM, elevated mDNAsi correlates with advanced tumor grade, worse overall survival (OS), and shorter progression-free survival (PFS), implicating both the primitive origin of malignant cells and the tumor TME as key determinants of aggressiveness [30]. Of the three biological EA clocks examined, ΔGrimAge2 and ΔDunedinPACE showed relatively few significant associations with tumor characteristics; however, ΔDunedinPACE was positively correlated with the astrocyte fraction. Genomic instability, characterized by aneuploidy and large-scale chromosomal alterations, is a recognized hallmark of both aging and cancer. Consistent with this, stemTOC was positively correlated with the fraction of the genome altered, whereas hypoClock showed a negative correlation with this measure. Microsatellite instability (MSI), while reported with low frequency in adult GBM, has been primarily linked to sporadic mismatch repair (MMR) loss rather than germline mutations, and its prognostic value in this context is limited [31]. Interestingly, a positive correlation between MSI-MANTIS scores and ΔDunedinPACE was observed (Figure. 1D).

Next, we utilized EA profiles to categorize GBM tumors into two distinct groups. Principal Component Analysis (PCA) was first performed on the EA matrix to reduce dimensionality and address multicollinearity among epigenetic clock features, followed by Linear Discriminant Analysis (LDA) to identify linear combinations that maximally separated the emerging groups. This approach classified GBM cases into two subgroups: EA-accelerated (EAA; n = 40), which exhibited accelerated aging across all biomarkers except hypoClock, and EA-decelerated (EAD; n = 69), characterized by elevated hypoClock scores alongside decelerated values for all remaining clocks (Figure. 1E).

### EA Stratifies GBM into Clinically Meaningful Subtypes

The distribution of assigned EA groups varied across established GBM subtypes, reflecting underlying differences in tumor biology and patient prognosis. Compared with the transcriptomic subtypes defined by Verhaak et al. [32], EAA cases were primarily associated with the classical (CL) subtype (64%), while mesenchymal (ME) and proneural (PN) subtypes constituted 16% and 20%, respectively. In contrast, EAD showed a greater proportion of the ME subtype (47.4%), with CL and PN subtypes accounted for 34.2% and 18.4 %, respectively (Figure. 2A). The CL subtype is often associated with *EGFR* amplification and a proliferative phenotype, whereas ME subtype is linked to increased invasiveness, upregulation of ME markers such as *CD44* and *CHI3L1*, and poorer prognosis [33]. Complementing the transcriptomic classification, we next examined the DNAm-based classification proposed by Ceccarelli et al.[34]. EAA tumors were mainly classified as LGm4 (84.2%), with smaller fractions of LGm5 (13.2%) and LGm6 (2.6%). Conversely, EAD tumors were predominantly LGm5 (66.2%), with fewer categorized as LGm4 (20.0%) and LGm6 (13.8 %) (Figure. 2B). LGm4 is associated with a more favorable prognosis and higher DNAm, aligning with CL-like profiles, whereas LGm5 indicates more aggressive tumor behavior, poorer survival, and overlaps with ME-like methylation patterns [34]. The DNAm status of the MGMT promoter also serves as an important metric for classifying GBM patients. It is a significant prognostic factor, correlating with improved OS and PFS, as well as a more favorable response to TMZ treatment [35, 36]. EAD tumors predominantly exhibited an unmethylated MGMT promoter (66.2%), whereas 53.8 % of EAA cases exhibited a methylated profile (Figure. 2C).

**Figure 2.**
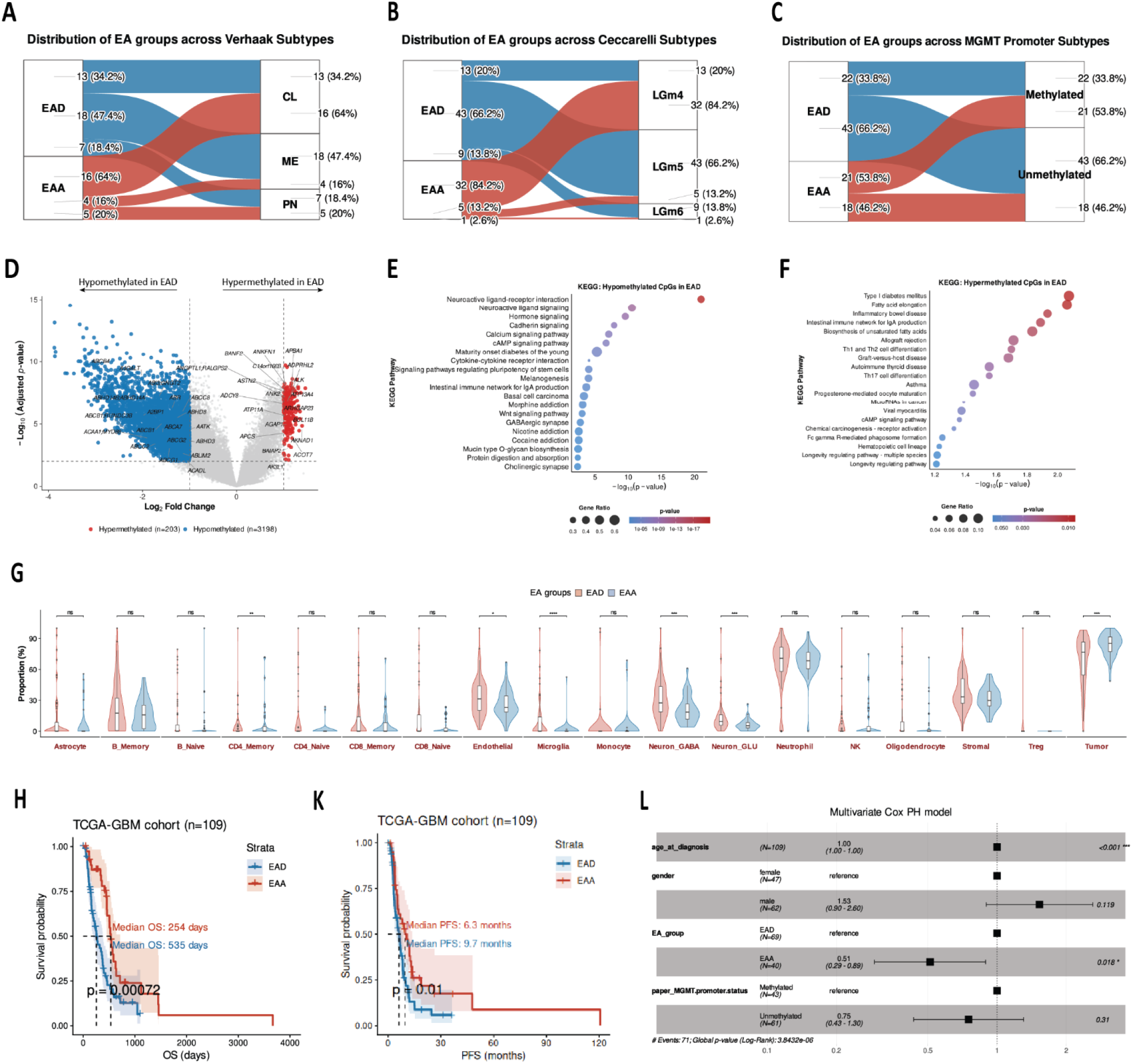
**(A-C)** Sankey plots depicting the proportional flows of EA groups to **(A)** Verhaak transcriptional subtypes, **(B)** Ceccarelli methylation subtypes, and **(C)** MGMT promoter status. **(D)** Volcano plot illustrating differentially methylated genes in EAD vs. EAA. Each dot represents a CpG probe, mapped to its corresponding gene. **(E)** KEGG overrepresentation enrichment analysis of hypomethylated probes in EAD. **(F)** KEGG overrepresentation analysis of hypermethylated probes in EAD. **(G)** Violin plots showing proportion of deconvoluted cell types in EAA, EAD groups. **(H)** Kaplan-Meier curve showing OS of EAA, EAD cases. **(K)** Kaplan-Meier curve showing PFS of EAA, EAD cases. Statistical significance assessed using the log-rank test (p-value shown on plot). **(L)** Forest plot illustrating multivariate Cox regression analysis in GBM patients, incorporating EA group as a covariate.

To compare DNAm profiles between EAA and EAD samples, we analyzed 337,202 CpG sites and identified 10,565 hypermethylated and 31,622 hypomethylated CpG sites in EAD relative to EAA (Figure. 2D, Supplementary material: Table S2). KEGG pathway enrichment analysis of differentially methylated probes (DMPs) showed hypomethylated probes significantly enriched in Neuroactive ligand-receptor interaction, Neuroactive ligand signaling, hormone, cadherin, and calcium signaling, and also pathways regulating pluripotency of stem cells (Figure. 2E). Hypermethylated probes were enriched in pathways such as Type I diabetes mellitus, Fatty acid elongation, and inflammatory bowel disease (Figure. 2F). We used a DNAm dataset from Drexler et al. [37] as an external validation cohort, comprising 209 EAD and 154 EAA cases (Supplementary material: Figure. S1A). In this dataset, hypomethylated probes in EAD were also enriched in neuroactive ligand-receptor interactions, cytokine-cytokine receptor interactions, and calcium signaling pathways (Supplementary material: Figure. S1B, C), while hypermethylated probes were mainly enriched in MAPK, Rap1, and Hippo signaling pathways (Supplementary material: Figure. S1D). Complete list of DMPs is provided in Supplementary material: Table S3.

Using the GIMiCC package [38], we deconvoluted the DNAm data across EA groups and compared the estimated proportions of eighteen cell types between EAA and EAD tumors. EAA GBMs exhibited a marked enrichment in malignant cell content, whereas EAD cases were significantly enriched for stromal cells, endothelial cells, memory CD4⁺ T cells, and brain-resident populations, including microglia, GABAergic neurons, and glutamatergic neurons (Figure. 2G). These findings suggest that EAA tumors are predominantly composed of malignant cells, whereas EAD tumors harbor a more complex TME enriched in stromal and neuronal constituents.

To compare the prognosis of the EA groups, we conducted Kaplan-Meier analysis, which revealed that EAD patients have notably poorer OS and PFS compared to EAA patients (Figure. 2H, K). The multivariate Cox proportional hazards model revealed that EA group is an independent predictor significantly associated with survival. Compared with EAD, patients in the EAA group showed a significantly decreased risk of the event, with a hazard ratio of 0.51 (95% CI: 0.29-0.89, p = 0.018), corresponding to an approximately 49% lower hazard (Figure. 2L). This indicates that membership in the EAD group is associated with worse survival outcomes relative to EAA, independent of other covariates. We replicated this finding in external validation cohort (Supplementary material: Figure. S1E).

### EA Subtypes exhibit a distinct TME architecture

Characterizing the link between EA profile and TME composition may uncover age-associated vulnerabilities in GBM, informing the development of age-personalized therapeutic strategies. To investigate expression modules associated with EA groups, we used an integrative analysis of paired bulk DNAm and RNA-seq datasets of GBM samples (n = 70). First, we computed a scale-free gene expression network (weighted correlation network analysis; WGCNA) resulting in gene expression modules, which were further correlated to EA groups through module significance measurement by quantifying the absolute correlation between the epigenetic signature and the individual module-derived gene expression profiles (Figure. 3A). We identified three expression modules significantly correlated with the EAA subtype: darkgreen (R² = -0.43, p = 0.004), grey60 (R² = -0.35, p = 0.02), and saddlebrown (R² = -0.41, p =0.006). Gene ontology (GO) enrichment analysis revealed that the darkgreen module was associated with regulation of growth factor receptor signaling and the ERK1/2 cascade, while grey60 was enriched for mitotic activity terms, and saddlebrown was associated with retinal and visual development gene sets. Four modules—black (R² = 0.34, p = 0.02), red (R² = 0.30, p = 0.04), turquoise (R² = 0.36, p = 0.018), and royalblue (R² = 0.31, p = 0.04)—were significantly correlated with the EAD subtype. The black module was associated with myelination and neuronal establishment, red was enriched for neuronal synaptic transmission and membrane potential terms, and turquoise was associated with leukocyte-mediated immunity. The royalblue module did not show significant enrichment for any GO term (Figure. 3B, C).

**Figure 3.**
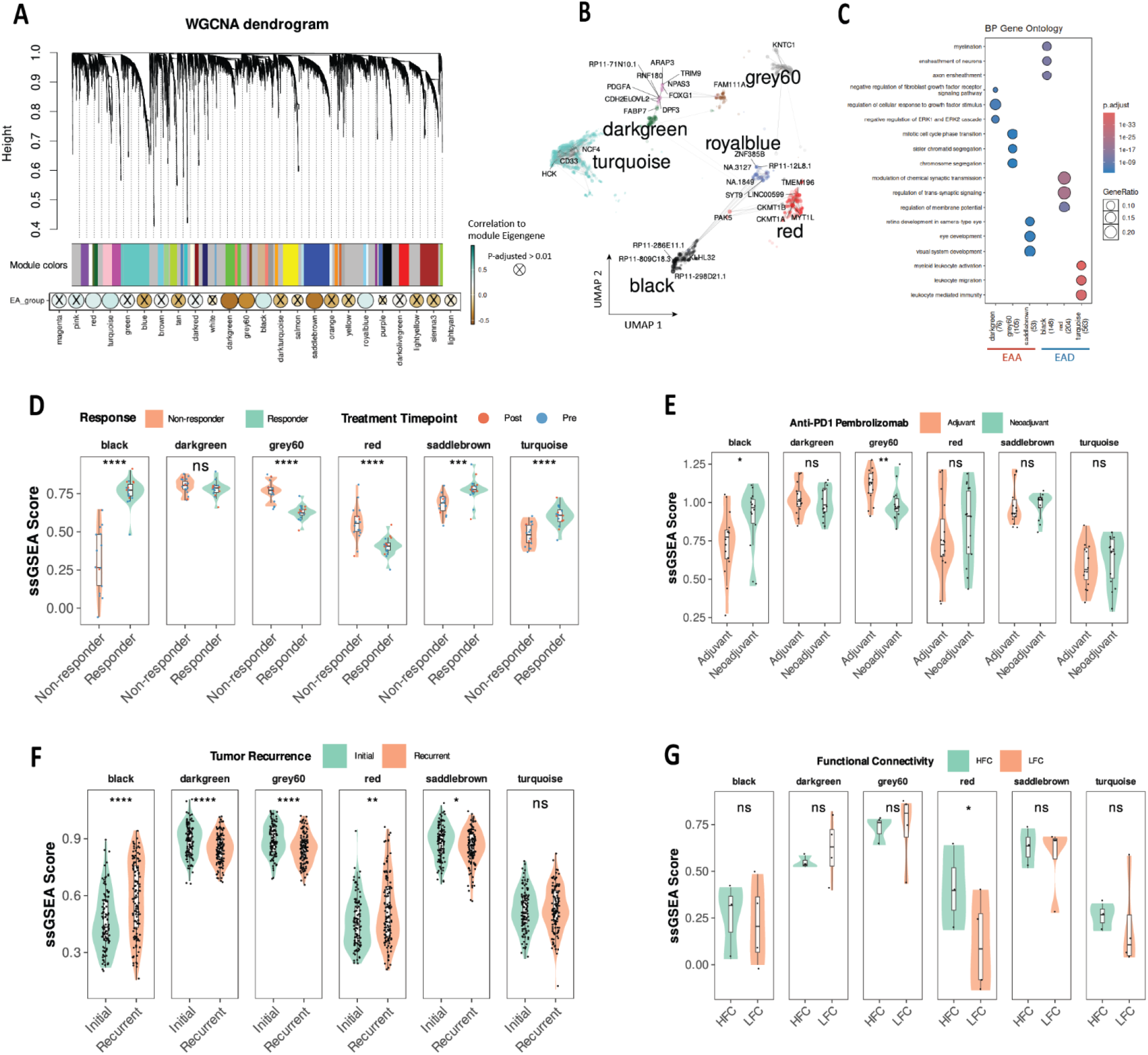
(**A**) Hierarchical dendrogram showing gene expression modules identified by WGCNA. The dot plot illustrates the Pearson correlation between identified modules and EA age, where dot size and color represent the correlation coefficient. Non-significant correlations are marked. (**B**) UMAP plot showing visualization of significantly correlated modules. (**C**) Gene ontology analysis of gene module eigengenes in EAA, and EAD. (**D**) Violin plots comparing WGCNA module eigengene ssGSEA enrichment scores between responder and non-responder GBM patients receiving anti-PD-1 therapy (nivolumab or pembrolizumab). (**E**) Violin plots comparing WGCNA module eigengene ssGSEA enrichment scores between GBM patients treated with adjuvant versus neoadjuvant pembrolizumab. (**F**) Violin plots comparing WGCNA module eigengene ssGSEA enrichment scores between initial and recurrent GBM tumor samples. (**G**) Violin plots comparing WGCNA module eigengene ssGSEA enrichment scores across HFC and LFC GBM tumors.

To evaluate the clinical relevance of the identified modules, we first used a longitudinal bulk RNA-seq dataset from GBM patients (n=17) treated with anti-PD-1 immunotherapy (nivolumab or pembrolizumab). This dataset originated from a study demonstrating that non-responder cases exhibit a distinct genomic profile linked to an immunosuppressive TME [39]. Significant enrichment of modules black, saddlebrown, and turquoise was observed in responder cases, while modules grey60, and red showed higher enrichment in non-responders (Figure. 3D). Subsequently, we compared module eigengene enrichment between neoadjuvant anti-PD-1 pretreated and adjuvant-treated recurrent GBM cohorts using their respective RNA-seq profiles [40]. The black module was significantly more enriched in the neoadjuvant group relative to the adjuvant group, whereas the grey60 module showed the opposite pattern, with greater enrichment in the adjuvant cohort (Figure. 3E).

Despite large-scale longitudinal studies investigating the evolutionary trajectories of recurrent GBMs, evidence for selective pressure against specific genetic alterations remains elusive. Investigating the dynamic interactions within the GBM TME during progression may therefore yield new therapeutic insights [41]. Accordingly, we analyzed paired initial and recurrent IDH-WT GBM samples (n=124) from the GLASS bulk RNA-seq dataset [41]. Modules darkgreen, grey60, and saddlebrown exhibited significantly higher enrichment in initial tumors. While turquoise showed no significant difference, black, and red modules were upregulated in recurrent tumors (Figure. 3F).

Additionally, we used a publicly available RNA-seq dataset from high functional (HFC, n=3) and low functional connectivity (LFC, n=4) GBMs and evaluated enrichment of module eigengenes. GBMs with HFC are characterized by increased integration of malignant cells into neural circuits and are associated with poorer survival and enrichment of immunosuppressive myeloid compartments [6, 42]. The red module showed a higher enrichment level in HFC compared to LFC (Figure. 3G).

In order to investigate EA subtypes at the single-cell level, we utilized the GBmap scRNAseq atlas [43]. To characterize the association of TME cell types with EA groups, we used the SCISSOR R package [44] and integrated the GBmap scRNA-seq atlas with TCGA bulk RNA-seq dataset (n=109) annotated for EA subtypes. Based on this analysis, EAA primarily correlated with malignant OPC-like, MES-like, and AC-like states. In contrast, EAD cases showed a positive association with oligodendrocytes, and malignant NPC-like states, as well as immune cell types such as TAMs, CD4/CD8 T-cells, and DCs (Figure. 4B).

**Figure 4.**
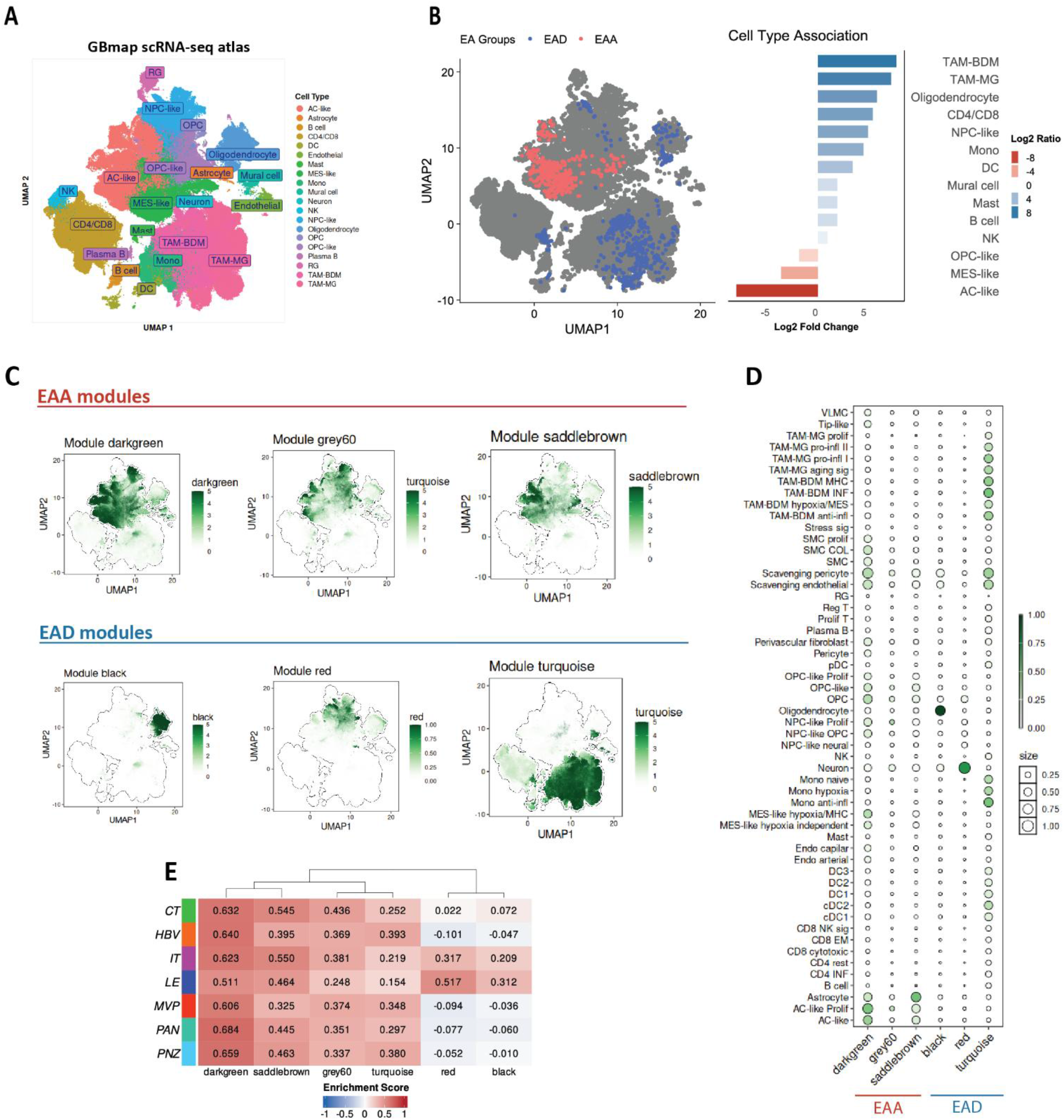
(**A**) UMAP dimensionality reduction of GBmap scRNAseq atlas. (**B**) UMAP plot showing the association of cell-type composition with EA groups. (**C**) Module eigengene expression of EAA, EAD GBM in GBmap scRNAseq atlas. (**D**) Heatmap dot plot showing expression enrichment of EAA, EAD-associated module eigengenes across GBmap scRNA-seq cell-type annotations. (**E**) Heatmap plot showing the scaled enrichment scores of identified module eigengenes across anatomical regions of GBM, using IVY-GAP bulk RNA-seq dataset.

We next projected the inferred module eigengene signatures onto the scRNA-seq UMAP. The darkgreen module was predominantly expressed in malignant cell states, as well as in normal astrocytes, oligodendrocyte progenitor cells (OPCs), brain pericytes, endothelial cells, and smooth muscle cells (SMCs). The grey60 module was highly expressed in malignant AC-like and NPC-like cells, whereas the saddlebrown module was expressed in non-malignant astrocytes and OPCs, along with malignant AC-like and OPC-like cells. The black and red modules were specifically enriched in normal oligodendrocytes and brain neurons, respectively. Finally, the turquoise module was expressed across immune clusters, particularly in subsets of TAMs, monocytes, and DCs, as well as in pericytes and endothelial cells (Figures. 4C and 4D).

In follow-up, we employed Ivy Glioblastoma Atlas Project (IVY-GAP) anatomical transcriptional atlas [45] to assess the expression scores of modules across anatomical regions (n=7) in GBM patients (n = 10). Cellular tumor (CT) region represents the dense core of proliferating tumor cells. Infiltrating tumor (IT) region consists of tumor cells that have invaded into surrounding normal brain tissue and leading edge (LE) is the tumor margin where tumor cells interact with normal brain parenchyma and may be less proliferative but more migratory. Importantly, microvascular proliferation (MVP) and hyperplastic blood vessel (HBV) regions are related to angiogenesis, immune response regulation, and wound healing. Pseudopalisading cells around necrosis (PAN), are characterized by hypoxia and stress responses. Modules red and black showed an enrichment pattern in IT, and LE. This finding aligns with a recent study demonstrating that malignant cells residing within infiltrated brain parenchyma show increased expression of neurodevelopmental pathways and synaptic genes [46]. While darkgreen, and saddlebrown showed a more ubiquitous anatomical expression, turquoise was largely enriched in HBV, MVP, and PNZ (Figure. 4E).

### Spatially resolved transcriptomics unveils module-specific expression landscapes and niche-niche interactome

Next, we utilized spatial transcriptomics (ST) data from Ravi et al. [47] to perform an in-depth examination of the TME architecture and cellular organization, with the goal of uncovering the spatial arrangement and relationships among EA-associated components (Figure. 5A, B).

**Figure 5.**
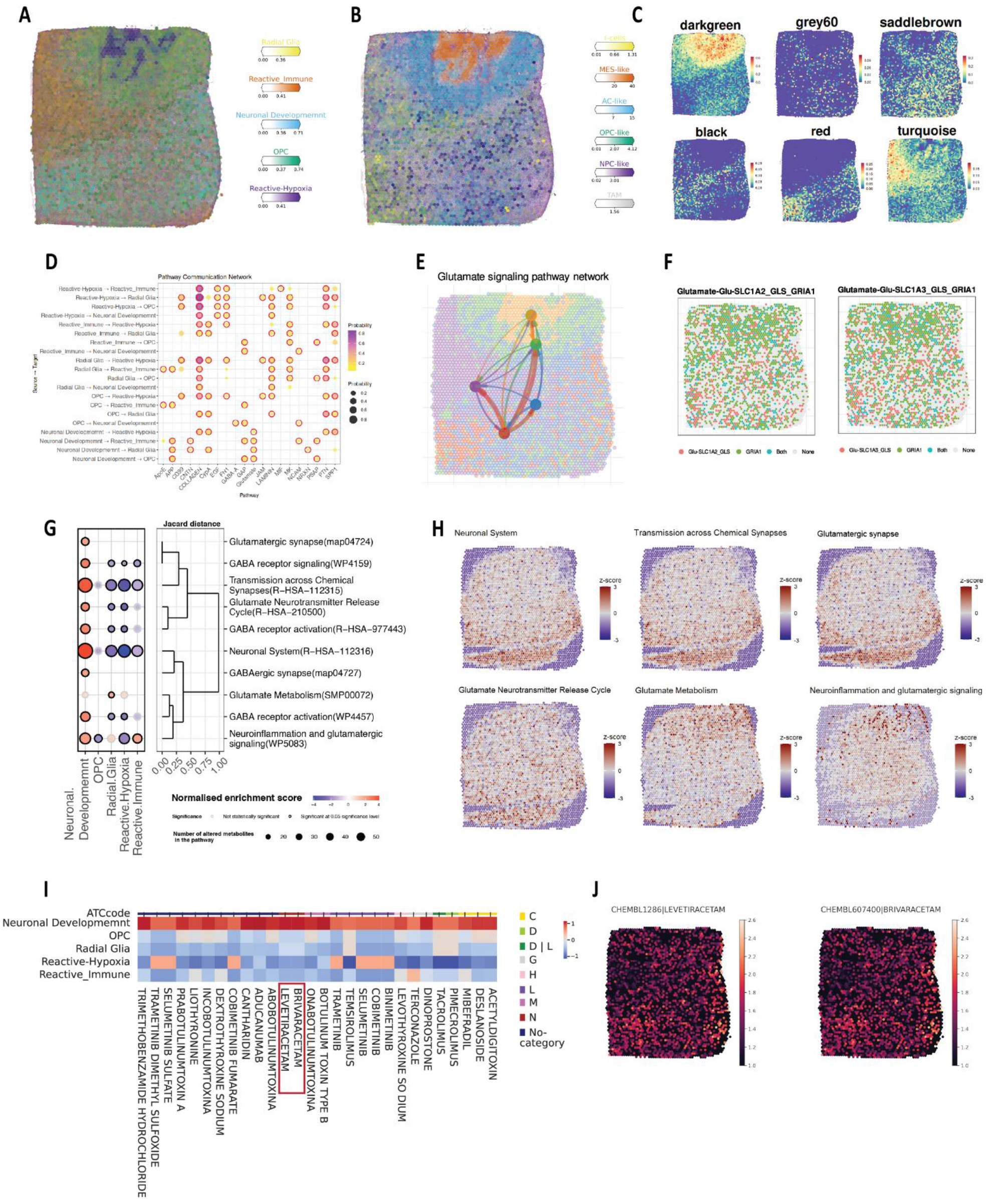
**(A)** Surface plot displaying the spatial enrichment scores of five transcriptional programs. **(B)** Surface plot showing the spatial deconvolution of cell types in Visium ST dataset from UKF275_T patient. **(C)** Surface plots showing spatial enrichment of module eigengenes. **(D)** Dot plot showing probability of inferred pathways between spatial niches. **(E)** Surface plot showing spatial direction of Glutamate signaling within tumor niches. **(F)** Spatial expression of top ligand-receptors of Glutamate signaling. **(G)** Heatmap dot plot showing differential multimodal pathway analysis across GBM spatial regions. **(H)** Spatial distribution of metabolic pathway enrichment scores derived from SM data, visualized as surface plots. **(I)** Heatmap plot of spatially-defined therapeutic enrichment analysis highlighting potential drugs targeting neural development regions. Rows represent GBM spatial transcriptional programs, and columns represent individual candidate drugs. Color scale reflects Drug2Cell enrichment scores, with red indicating higher scores and blue indicating lower scores. Drugs are grouped and color-coded along the top bar according to their Anatomical Therapeutic Chemical (ATC) classification: C, Cardiovascular system; D, Dermatological; D|L, Dermatological/Antineoplastic and immunomodulating agents; G, Genito-urinary system and sex hormones; H, Systemic hormonal preparations; L, Antineoplastic and immunomodulating agents; M, Musculoskeletal system; N, Nervous system; No-category, drugs without an assigned ATC category. **(J)** Surface plots illustrating the predicted spatial distribution of drug targeting scores for LEV (left) and BRV (right).

Spatial expression analysis of module eigengenes revealed distinct associations with specific ST programs. The turquoise module was predominantly expressed in the reactive immune niche, characterized by high abundance of immune cell populations, including TAMs and T cells. In contrast, the red module was significantly enriched in the neuronal development niche, marked by the presence of neurons, and NPC-like cells. The darkgreen module showed strong enrichment in the hypoxia-reactive region, while the black and grey60 modules were enriched in the OPC niche surrounding this region. The saddlebrown module was broadly expressed across the OPC region (Figure. 5C).

Because the EAD-associated red module was enriched in the neuronal-development region, we focused on this compartment and inferred its spatial communication networks with immune and malignant niches using the CellChat package [48], aiming to identify key pathways through which neurons and GSCs promote immunosuppression and tumor recurrence. Figure. 5D shows the top 20 most probable inferred pathways. Notable active pathways connecting the neuronal development niche and the immune-reactive region included Amyloid Precursor Protein (APP), Contactin (CNTN), Growth-Associated Protein 43 (GAP43),

Neural Cell Adhesion Molecule (NCAM), and Prosaposin (PSAP). Additionally, the neuronal development niche was found to interact with the reactive-hypoxia niche via the COLLAGEN, Cyclophilin A (CypA), Pleiotrophin (PTN), and Secreted Phosphoprotein 1 (SPP1) signaling pathways. Notably, glutamate signaling emerged as a dominant axis, active in both neuronal-immune and neuronal-hypoxic interactions, suggestive of a potential dual role in immunomodulation and adaptive survival via glioma-neuron-immune crosstalk. GBM cells release excessive glutamate into the tumor microenvironment, driving excitotoxicity and promoting tumor growth [49]. This glutamatergic signaling contributes to metabolic reprogramming, transcriptional switching, and the formation of synaptic-like connections within the TME [50]. At the cellular level, glutamate activates calcium-permeable AMPA receptors on GSCs, stimulating their proliferation and invasive capacity [51]. Among metabotropic receptors, metabotropic glutamate receptor 3 (GRM3) is predominantly expressed in GBM, where it regulates GSC self-renewal [52]. Beyond direct effects on tumor cells, GBM regions with enhanced neuronal connectivity exhibit regional immunosuppression characterized by anti-inflammatory tumor-associated macrophages (TAMs), and pharmacological glutamate inhibition shifts TAMs toward less immunosuppressive states [6].

To further investigate the neuronal development regions, we analyzed ST-paired, spatially resolved metabolomics (SM) data generated by MALDI Fourier-transform ion cyclotron resonance imaging mass spectrometry (MALDI-FTICR-MSI) [47]. Differential multimodal pathway analysis integrating ST and SM data revealed significant enrichment of Reactome pathways associated with the neuronal system, transmission across chemical synapses, neuroinflammation, and glutamatergic signaling within the neuronal development region (Figure. 5G). Moreover, SM-based metabolite activity analysis showed that glutamate metabolism and signaling were most elevated in the reactive hypoxia and neuronal development niches, whereas pathways linked to neuronal function and activity were highest specifically within the neuronal development niche (Figure. 5H).

To identify potential therapeutic agents targeting the neuronal development niche and disrupt neuronal-immune crosstalk, we applied the Drug2Cell algorithm [53] within the ST framework. Leveraging pharmacogenomic databases (e.g., ChEMBL), Drug2Cell prioritizes drug candidates based on their expression profiles, offering potential efficacy in modulating niche-specific cellular populations. The results pointed to several members of the botulinum neurotoxin (BoNT) family. Additionally, the antiepileptic medications LEV and brivaracetam (BRV) were among the suggested drugs with the highest specificity (Figure. 5I, J). LEV reduces excessive glutamate signaling by presynaptic inhibition of glutamate release via SV2A binding and modulation of P/Q-type calcium channels, dampening excitatory neurotransmission and protecting neural circuits from excitotoxicity. This mechanism underlies its efficacy in epilepsy and potentially other neurological disorders involving glutamate dysregulation such as GBM [54]. Several large retrospective studies and meta-analyses report that LEV use is associated with improved OS and PFS in GBM patients [55]. Mechanistically, LEV may enhance TMZ efficacy by modulating *MGMT* expression and inhibiting immunosuppressive microglial polarization [56]. A double-blind randomized trial is currently underway to assess LEV’s impact on GBM patient outcomes [57]. RIL is another anti-glutamatergic agent with well-established neuroprotective properties against excitotoxicity. It was selected alongside the Drug2Cell-identified LEV to introduce mechanistic diversity into the experimental design. RIL exerts its effects through multiple complementary pathways: blocking voltage-gated sodium channels to suppress presynaptic glutamate release, enhancing astrocytic glutamate reuptake, and modulating postsynaptic NMDA and kainate receptors [36, 58]. Including both drugs enables the discrimination of mechanism-specific effects from those shared across the anti-glutamatergic drug class.

### Individual and combined effects of LEV, RIL, and TMZ on cell viability

The anticancer effects of LEV and RIL were evaluated individually and in combination with TMZ in the U87MG and A172 GBM cell lines, as well as in the non-cancerous human fetal foreskin fibroblast (HFFF2) cell line, using the MTT assay at 24- and 48-hours post-treatment. Our results demonstrated that LEV, RIL, and TMZ induced dose- and time-dependent cytotoxicity across all tested cell lines. After 24 hours of exposure, the half-maximal inhibitory concentration (IC₅₀) values for LEV were 1113 µg/mL in U87MG cells (Figure. 6A), 1430 µg/mL in A172 cells (Figure. 6B), and 1841 µg/mL in HFFF2 cells (Figure. 6C). For RIL, the corresponding 24-hour IC₅₀ values were 27.97 µg/mL in U87MG cells (Figure. 6D), 24.07 µg/mL in A172 cells (Figure. 6E), and 39.57 µg/mL in HFFF2 cells (Figure. 6F). Similarly, the 24-hour IC₅₀ values for TMZ were 56.77 µg/mL in U87MG cells (Figure. 6G), 116.4 µg/mL in A172 cells (Figure. 6H), and 34.77 µg/mL in HFFF2 cells (Figure. 6K). Notably, the cytotoxic effects of both LEV and RIL were significantly enhanced following 48 hours of treatment. A summary table of IC₅₀ values is shown in Figure. 6J.

**Figure 6.**
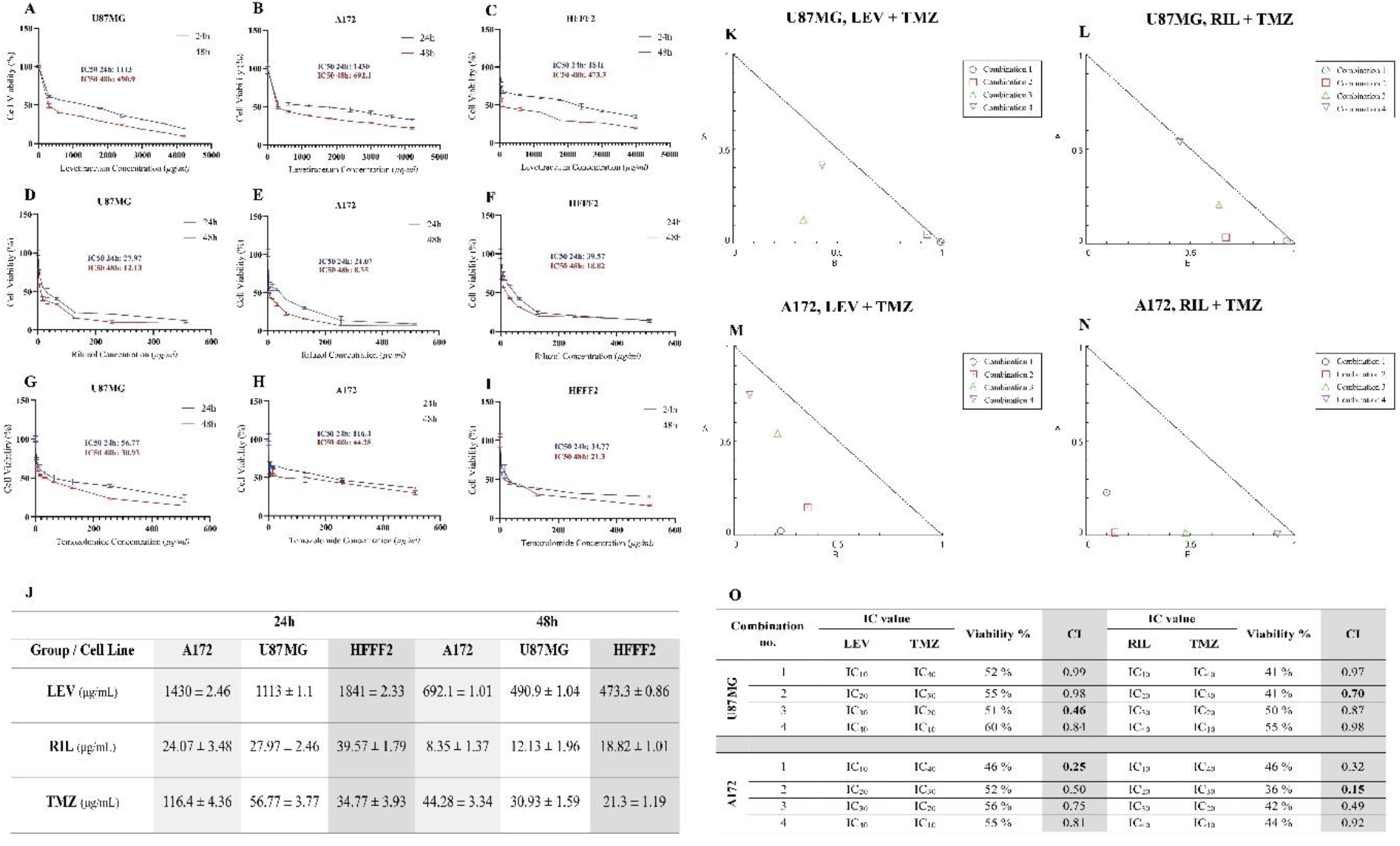
Cytotoxic and synergistic profiles of LEV, RIL, and TMZ on GBM and normal cell lines. **(A-I)** Dose and time-dependent cell viability curves of U87MG, A172, and HFFF2 cells treated with individual compounds for 24 and 48 hours, evaluated by MTT assay **(J)** Summary table of IC₅₀ values (µg/mL). **(K-N)** Isobologram analysis and CI plots demonstrating synergistic interactions of LEV+TMZ and RIL+TMZ combinations in U87MG and A172 lines. (**O**) Summary table of combination values. Data is presented as the mean ± SD of three independent experiments. (n = 3).

Combination treatments of LEV and RIL with TMZ showed synergistic effects in both GBM cell lines. In U87MG cells, LEV combination 3 (LEV at IC30 with TMZ at IC20; CI=0.46) and RIL combination 2 (RIL at IC20 with TMZ at IC30; CI=0.70) demonstrated the most pronounced synergy (Figure 6K, L). Similarly, in A172 cells, LEV combination 1 (LEV at IC10 with TMZ at IC40; CI=0.25) and RIL combination 2 (RIL at IC20 with TMZ at IC30; CI=0.15) exhibited synergy, which was confirmed by isobologram analysis (Figure 6M, N). A summary table of combination index values is presented in Figure. 6O.

### LEV and RIL, in combination with TMZ, modulate mRNA levels of stemness, neuronal-activity, and immunosuppression marker genes in vitro

We evaluated transcriptional responses associated with stemness, neuronal activity, and immunosuppression in U87MG and A172 GBM cells following treatment with LEV, RIL, and TMZ individually, as well as in combination with TMZ.

In U87MG cells, TMZ monotherapy modestly increased the expression of the stemness regulators *SOX2* and *EZH2* by approximately 2.8-fold and 5-fold, respectively, whereas in A172 cells, TMZ induced a 1.8-fold increase in *SOX2* and a more pronounced 2.1-fold upregulation of *EZH2* (Figure. 7A). TMZ paradoxically promotes cellular plasticity by increasing the frequency of GSCs and enabling differentiated tumor cells to reprogram into stem-like states [59]. Conversely, TMZ has also been reported to preferentially deplete GSCs, particularly in MGMT-negative tumors [60]. LEV treatment induced significant downregulation of *SOX2* in both cell lines (0.44–fold in U87MG and 0.36–fold in A172), while causing only a mild, non–significant reduction in *EZH2* expression. The combination of LEV with TMZ did not result in any significant changes in the expression of *SOX2* or *EZH2*. RIL treatment significantly downregulated *SOX2* expression by 0.55–fold in U87MG cells. Similarly, *EZH2* expression decreased by 0.6–fold in U87MG and 0.4–fold in A172 following RIL treatment. The combination of TMZ with RIL caused a mild, non–significant upregulation of both genes compared with TMZ monotherapy (Figure. 7A).

**Figure 7.**
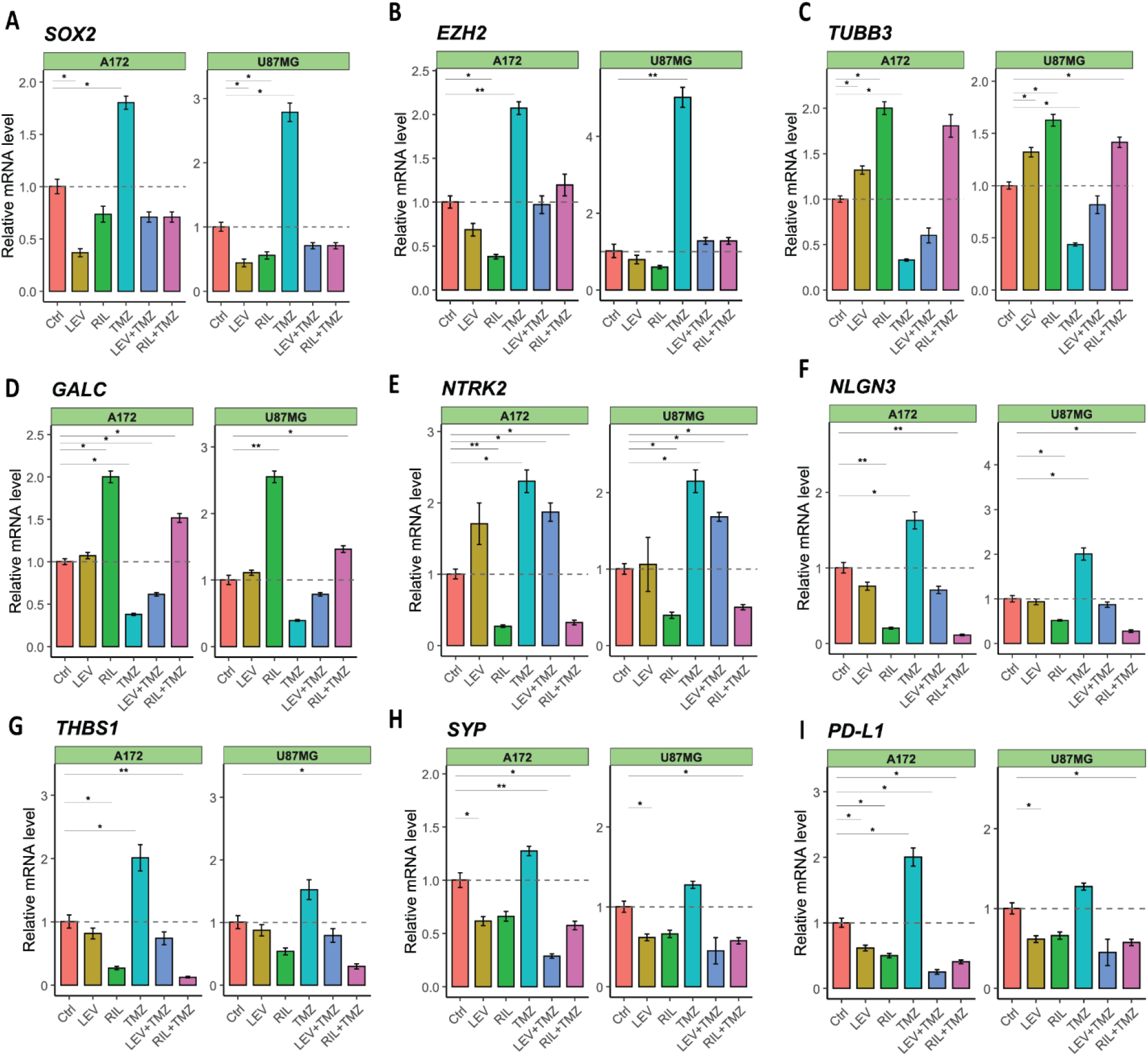
**(A)** Bar plots showing RT-qPCR relative mRNA levels of *SOX2* **(A)**, *EZH2* **(B)**, *TUBB3* **(C)**, *GALC* **(D)**, *NTRK2* **(E)**, *NLGN3* **(F)**, *THBS1* **(G)**, *SYP* **(H)**, and *PD-L1* **(I)** in A172 and U87MG cells under Ctrl, LEV, RIL, TMZ, and combination treatments (LEV+TMZ, RIL+TMZ). Expression levels were normalized to *GAPDH* internal housekeeping gene and are presented relative to the control (dashed line, set to 1). Bars represent mean ± SEM from independent experiments. Statistical significance between groups is indicated as p < 0.05 (*), p < 0.01 (**), p < 0.001 (***).

In contrast to stemness-associated genes, the neuronal differentiation markers *TUBB3* and *GALC* were downregulated following TMZ monotherapy in U87MG cells, with expression reduced by approximately 0.45-fold and 0.4-fold, respectively. Similarly, in A172 cells, TMZ treatment resulted in a modest 0.3-fold reduction in *TUBB3* and a 0.4-fold decrease in *GALC* expression. LEV treatment significantly upregulated *TUBB3* expression in both cell lines; but did not significantly alter *GALC* expression. Its combination with TMZ led to decreased expression of *GALC* by 0.8-fold in U87MG and 0.6-fold in A172 cells, and of *GALC* by 0.8-fold and 0.6-fold, respectively. RIL treatment markedly upregulated *TUBB3* expression, with approximately 1.6-fold and 2-fold increases in U87MG and A172 cells, respectively. Likewise, *GALC* expression was elevated following RIL treatment, showing 2.55-fold upregulation in U87MG and 2-fold in A172 cells. Notably, combining TMZ and RIL was most effective in restoring glial identity and promoting neuronal differentiation. Specifically, this treatment triggered a ∼1.4-fold upregulation of *TUBB3* in U87MG cells and a ∼1.8-fold increase in A172 cells. Similarly, *GALC* expression increased by 1.4-fold in U87MG cells and 1.5-fold in A172 cells (Figure. 7C, D).

Treatment with TMZ monotherapy resulted in the upregulation of key neuron-glioma communication mediators in both U87MG and A172 glioma cell lines. In U87MG cells, TMZ treatment led to a doubling in *NTRK2* and *NLGN3* expression, and a 1.2-fold increase in *SYP*. Similarly, in A172 cells, TMZ increased the expression of *NTRK2*, *NLGN3*, and *SYP* by 2.3-fold, 1.6-fold, and 1.3-fold, respectively. *THBS1* showed a modest insignificant upregulation in U87MG and a 2-fold upregulation in A172 after TMZ treatment. These findings align with previous findings, showing that persistent TMZ pressure can select for subclones with altered metabolism that enhance glutamate release, supporting survival and excitotoxic damage to nearby neurons [49]. In contrast, the addition of LEV or RIL reversed this effect. While LEV monotherapy had no significant impact on the expression of *NTRK2, NLGN3*, and *THBS1*, it significantly downregulated *SYP* levels by approximately 0.6-fold in both U87MG and A172 cells. The combination of LEV and TMZ resulted in a marked downregulation of *NTRK2* by 0.87-fold in U87MG cells and 0.7-fold in A172 cells. Although this combination caused a non-significant downregulation of *NLGN3* and *THBS1*, it significantly downregulated *SYP* levels by 0.45-fold and 0.3-fold in U87MG and A172 cells, respectively. RIL monotherapy significantly reduced the expression of *NTRK2* (0.4-fold), *NLGN3* (0.5-fold), and *THBS1* (0.5-fold), while having no significant effect on *SYP* expression. The combination of RIL with TMZ resulted in the downregulation of *NTRK2* (0.5-fold), *NLGN3* (0.3-fold), *THBS1* (0.3-fold), and *SYP* (0.6-fold) in U87MG cells. Similarly, in A172 cells, the combination treatment downregulated mRNA levels of *NTRK2* (0.3-fold), *NLGN3* (0.1-fold), *THBS1* (0.1-fold), and *SYP* (0.6-fold) (Figure. 7E-H). *NTRK2* encodes TrkB, the receptor for brain-derived neurotrophic factor (BDNF), an activity-dependent neurotrophin implicated in glioma progression [61]. Disruption of BDNF-TrkB signaling has been shown to suppress tumor growth and confer a survival benefit in mouse models [5]. TrkB inhibition has also been shown to significantly reduce the viability of both U87MG and A172 GBM cell lines [62]. NLGN3 is an activity-dependent synaptic adhesion molecule cleaved and secreted by neurons and oligodendrocyte precursor cells in the GBM TME via ADAM10. Secreted NLGN3 acts as a potent mitogen promoting tumor proliferation, migration, and invasion, and its elevated levels are associated with poor survival and recurrence. Accordingly, ADAM10 inhibition has emerged as a promising strategy to block NLGN3 secretion and suppress GBM progression [4, 61]. As a secreted synaptogenic factor, *THBS1* promotes neuron-glioma synapse formation and stability in HFC GBM regions, thereby enhancing glutamatergic coupling and tumor proliferation [6].

TMZ monotherapy induced a significant upregulation of PD-L1 (*CD274*) expression, resulting in a 2-fold increase in A172 cells, whereas only a mild, non-significant increase was observed in U87MG cells. This observation aligns with previous reports indicating that TMZ can promote immune escape in GBM cells by upregulating PD-L1 [63]. LEV monotherapy resulted in a 0.6-fold downregulation of PD-L1 in both cell lines, while RIL significantly decreased its expression to 0.5-fold in A172 cells. Combining TMZ with either LEV or RIL attenuated the TMZ-induced increase in PD-L1 expression. Specifically, the LEV and TMZ combination reduced PD-L1 expression to 0.25-fold in A172 cells, while RIL and TMZ combination reduced levels to 0.5-fold and 0.4-fold in U87MG and A172 cells, respectively (Figure. 7I).

### Individual and combined effects of LEV, RIL, and TMZ on apoptosis

To assess the effects of LEV, RIL, and TMZ on apoptosis, Annexin V-FITC/propidium iodide (PI) double staining followed by flow cytometric analysis was performed in U87MG and A172 cells following treatment. In U87MG, TMZ monotherapy resulted in 13.7% early apoptotic, 6.17% late apoptotic, and ∼20% total apoptotic cells (P < 0.001). LEV monotherapy induced 14% early apoptotic, 5.37% late apoptotic, and 19.3% total apoptotic cells, while RIL induced 8.2% early apoptotic, 4.05% late apoptotic, and ∼12% total apoptotic cells. Combining either LEV or RIL with TMZ markedly increased cell death. Specifically, the LEV+TM Z combination yielded 20.3% early apoptotic, 8.3% late apoptotic, and ∼28% total apoptotic cells. Similarly, the RIL+TMZ combination resulted in 16.33% early apoptotic, 10% late apoptotic, and 26.3% total apoptotic cells (Figure. 8A, B).

**Figure. 8.**
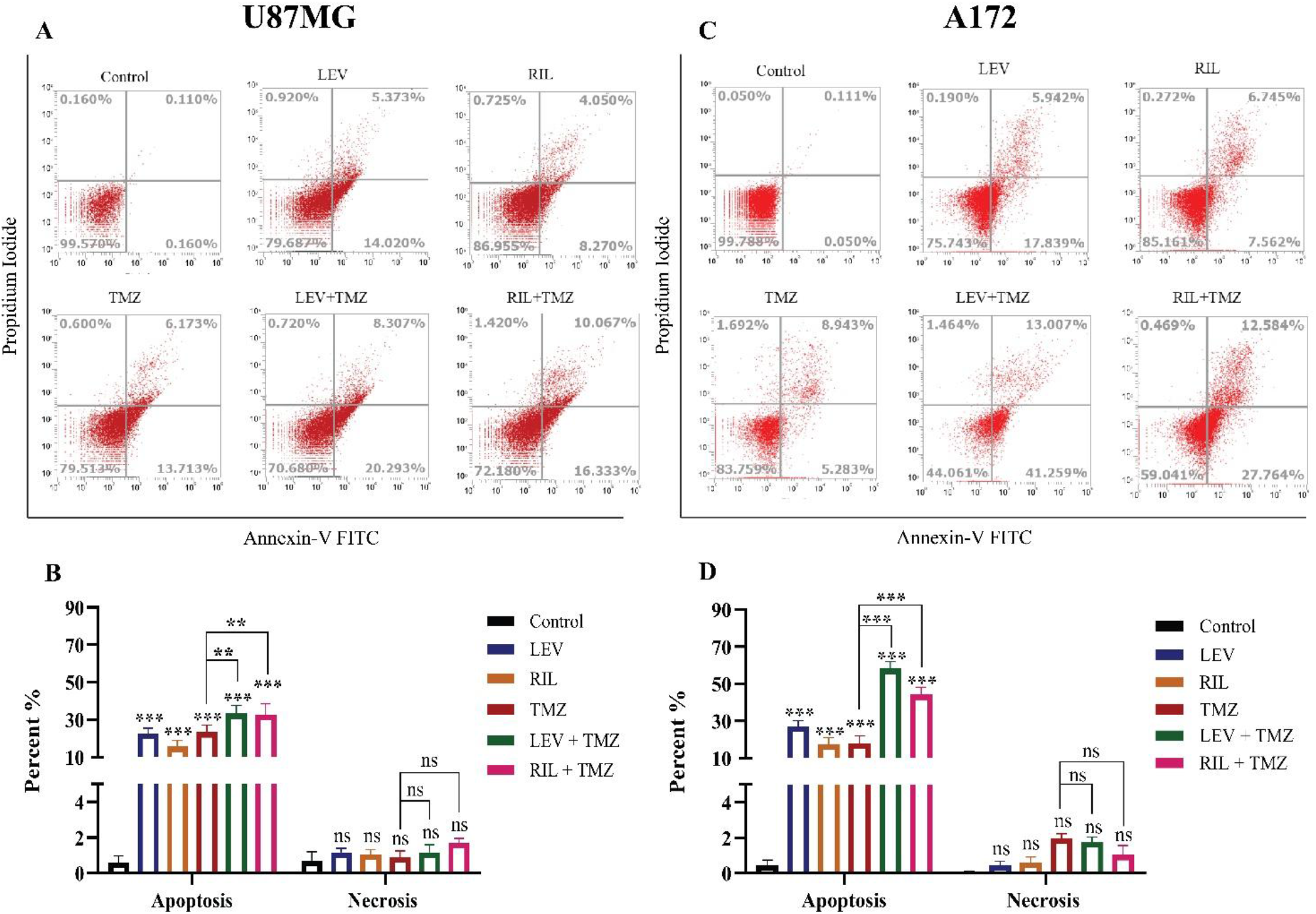
Flow cytometry-based apoptosis analysis of GBM cell lines after 24h treatment with LEV, RIL, and TMZ. Representative dot plots of Annexin V-FITC versus propidium iodide (PI) staining are shown for (**A, B**) U87MG and (**C, D**) A172 cells treated with vehicle Ctrl, TMZ, LEV, RIL, and their combinations (LEV+TMZ and RIL+TMZ). Quadrants represent viable cells (Annexin V⁻/PI⁻), early apoptotic cells (Annexin V⁺/PI⁻), late apoptotic cells (Annexin V⁺/PI⁺), and necrotic cells (Annexin V⁻/PI⁺). Quantitative analysis of apoptotic and necrotic populations is summarized in bar graphs for (**B**) U87MG and (**D**) A172 cells. Data are presented as mean ± SD from at least three independent experiments. Statistical significance was determined relative to control or between indicated groups (*p < 0.05, **p < 0.01, ***p < 0.001; ns, not significant).

In A172 cells, treatment with TMZ alone induced apoptosis in 14.22% of cells (5.28% early and 8.94% late apoptosis), while LEV monotherapy resulted in 23.77% apoptosis (17.83% early and 5.94% late) and RIL monotherapy yielded 14.3% apoptosis (7.56% early and 6.74% late). The addition of either LEV or RIL to TMZ significantly augmented apoptotic rates. Specifically, the LEV+TMZ combination increased total apoptosis to 54.25% (41.25% early and 13% late), whereas the RIL+TMZ combination elicited a more pronounced effect, achieving 40.34% total apoptosis (12.58% early and 27.76% late). Notably, apoptosis induced by LEV+TMZ was significantly greater than that observed with any single-agent treatment (Figure. 8C, D).

## Methods

### Analytical Methods

#### Computation of EA clocks and EA acceleration

We calculated three classes of DNAm-based aging biomarkers: chronological (Horvath1, AltumAge, and DNAmClockCortical), biological (HRSInChPhenoAge, GrimAge2, and DunedinPACE), and mitotic (epiTOC2, stemTOC, and HypoClock), using the MethylCIPHER R package [64] and the pyaging Python package.[65]. The EAA metric was determined solely for biological and chronological clocks, not for mitotic clocks. To calculate this acceleration, we subtracted CA from EA:

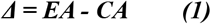

Mitotic clocks were excluded from EAA calculations because they primarily reflect cellular replication history and replicative senescence rather than true biological aging, rendering acceleration metrics derived from them not meaningful.

#### Reanalysis of publicly available bulk RNA-seq datasets from human glioma

Processed bulk RNA-seq count matrices and associated clinical annotations for the TCGA-GBM cohort was retrieved from cBioPortal (https://www.cbioportal.org/) using the TCGAbiolinks R package (v3.19). Datasets were subsequently filtered to retain only primary tumor samples, yielding 236 IDH-WT cases.

Gene expression data from the GLASS consortium were accessed via Synapse (https://www.synapse.org/glass). The downloaded file (“transcript_count_matrix_all_samples.tsv”) contained read counts at the splice isoform level, which were aggregated by summing across isoforms for each gene (“gene_id”) and rounded to the nearest integer to obtain gene-level count matrices. Replicate samples were excluded prior to analysis, and for cases with multiple recurrences, only the most recent recurrent sample was retained. PCA was performed on the top 1,000 most highly variable genes across all samples. Three samples were identified as outliers and removed from their respective batches: GLSS-SM-R099-R1-01R-RNA-MNTPMI, GLSS-SM-R111-R1-01R-RNA-WM5ESA, and GLSS-CU-R004-TP-01R-RNA-U0DEP1.

Raw FASTQ files (n = 32) from Zhao et al. [39] were first evaluated for quality using FastQC (v0.12.1; https://github.com/s-andrews/FastQC) with default parameters. Adapter sequences and low-quality bases were subsequently trimmed using TrimGalore (v0.6.10; https://github.com/FelixKrueger/TrimGalore) under default settings. Trimmed reads were aligned to the human reference genome (hg38) and quantified at the transcript level using Salmon [66]. A gene-by-sample count matrix was then constructed from the resulting alignments and subjected to variance-stabilizing transformation (VST). PCA was conducted on the transformed expression values using the top 1,000 most variable genes, and results were visualized accordingly. Donor-associated batch effects were corrected using the removeBatchEffect function from the limma package (RRID:SCR_010943, v3.36.5).

Bulk RNA-seq data were obtained from the IVY-GAP, encompassing 279 tumor fragment samples derived from two cohorts: 122 samples from 10 tumors comprising the anatomic structure cohort, and 157 samples from 34 tumors comprising the cancer stem cell cohort [45]. Normalized expression counts were stratified by tumor location, and location-specific expression patterns were visualized as heatmaps generated using ComplexHeatmap (v3.19).

#### Integrative analysis of DNAm and RNA-seq

Co-expression networks were constructed using the WGCNA R package (v0.2.2) [67], with EA labels incorporated as phenotypic traits. Network topology analysis identified an optimal soft-thresholding power of 12 to satisfy scale-free topology. Module eigengenes were projected into a reduced dimensions space using UMAP (*ModuleUMAPPlot*). Functional enrichment of the top 100 module genes was assessed via Gene Ontology analysis (*compareCluster*), with pathway activation patterns visualized using clusterProfiler.

#### Survival analysis

We performed survival analysis using both Kaplan-Meier estimation and Cox proportional hazards (CoxPH) regression. The survival R package (v3.6) was used to fit multivariate CoxPH models, assessing the association between EA groups and survival outcomes.

#### Correlation of phenotypic and genomic features

Scores for genomic features (fraction of genome altered, aneuploidy, microsatellite instability (MSI-MANTIS and MSI Sensor), and mutational burden) were downloaded from cBioPortal from PanCancerAtlas TCGA-GBM [68]. Tumor stemness scores (mRNAsi and mDNAsi) were retrieved as computed by Malta et al. [30].

#### SCISSOR analysis

We performed integrative analysis of bulk RNA-seq and single-cell RNA-seq data using SCISSOR (v2.1.0) [44], modeling EA groups as the dependent variable in a logistic regression framework.

#### Analysis of ST data

Using Scanpy (v1.11.3) in Python, we log-normalized the ST datasets [47] by scaling counts per spot to 10,000 and applying log1p transformation. The top 5,000 highly variable genes were selected (Seurat method), followed by dimensionality reduction, neighborhood graph construction, and UMAP projection—each slide was processed independently. For spatial cell-type mapping, we applied cell2location (v0.1.3) [69], a Bayesian tool that decomposes ST data using scRNA-seq-derived signatures. First, it models cell-type profiles via negative binomial regression, then performs non-negative factorization to estimate spatially resolved cell-type abundances. Spatial ligand-receptor interaction analysis was conducted using CellChat (version 1.5.0) [48].

#### Drug prediction for TME

To identify candidate drug repurposing opportunities targeting the neuronal-developmental niche, we leveraged ST data and applied Drug2Cell [53], an enrichment-based framework, to predict compounds with high specificity for this niche.

#### Analysis of SM data

After initial pre-processing, imzML object was loaded using *readImzML()* of Cardinal package v.3.10. MALDI samples were examined individually via the R package SpaMTP version 1.0 [70]. Mass peaks were binned at a 250-ppm resolution, producing 5411 detectable m/z features. Sample annotations were performed against the Lipid Maps database using SpaMTP’s *AnnotateSM()* function. Principal component analysis (PCA) was performed, with dimensionality reduction applied to each sample based on the leading 30 principal components. Pseudo-bulking differential metabolite abundance was assessed cluster-wise through the SpaMTP *FindAllDEMs()* function. The top 10 m/z values from each cluster across all samples were then compiled, and hierarchical clustering was applied to group akin clusters by their pseudo-bulked expression patterns. Additional differential abundance analysis identified metabolites with significant enrichment in neuronal-development clusters versus others in each sample.

#### Cell state composition analysis

Cell type deconvolution of bulk DNAm data from glioma samples was performed using the Glioma Immune Microenvironment Composition Calculator (GIMiCC) package [38], a computational tool specifically developed to estimate the proportions of multiple cell types within glioma tissues based on DNAm profiles.

#### Differential methylation analysis

Initially, functionally normalized methylation values were transformed into M-values with the *getM()* function from the minfi package (v1.54.1), while negative control probes were retrieved using the *getINCs()* function from MissMethyl package [71]. Probe annotations were obtained using the minfi package, using *getAnnotation()* function and analyses were restricted to promoter–associated probes defined as CpGs mapped to TSS1500/TSS200, 5’UTR, and first exon. Differentially methylated probes between groups were identified using linear models fitted to M values with the limma package v.3.66, including covariates such as age, sex. Resulting P values were corrected for multiple testing using the Benjamini-Hochberg false discovery rate procedure, and probes with an FDR < 0.01 and an absolute delta–beta ≥ 0.5 were considered significantly differentially methylated.

### Experimental Methods

#### Culture of human GBM cell lines

U87MG (ATCC® HTB-14™) and A172 (ATCC® CRL-1620™) GBM cell lines were cultured in RPMI supplemented with 10% fetal bovine serum (FBS) and 1% penicillin-streptomycin at 37°C with 5% CO₂. Cells were passaged at 80-90% confluency using 0.25% trypsin-EDTA, neutralized with complete medium, centrifuged (300 × g, 5 min), and reseeded at 5,000-15,000 cells/cm² (U87MG: 5,000-10,000; A172: 10,000-15,000). Mycoplasma testing was performed monthly, and cell morphology was routinely verified by microscopy. For cryopreservation, cells were suspended in 90% FBS/10% DMSO, frozen at -80°C, and stored in liquid nitrogen.

#### Cell-viability assay

Cell viability was determined using the MTT colorimetric assay, which quantifies the reduction of 3-(4,5-dimethylthiazol-2-yl)-2,5–diphenyltetrazolium bromide to insoluble formazan by metabolically active cells. U87MG and A172 cells were seeded in 96–well plates at a density of 1.0 × 10³ cells per well in 200 µL of RPMI–1640 medium supplemented with 10% FBS; the non–malignant HFFF2 fibroblast cell line was included to evaluate the safety profile of RIL, LEV, and TMZ. Following a 24-hour attachment period, cells were exposed to increasing concentrations of LEV (300-4200 µg/mL) and RIL (4-512 µg/mL), and TMZ (4-512 µg/mL) using eight concentration levels for each compound. At 24-and 48-hours post–treatment, the culture medium was replaced with 100 µL of fresh medium and 50 µL of MTT solution (5 mg/mL, Sigma–Aldrich) and incubated for 4 hours at 37°C to allow formazan crystal formation. The supernatant was then carefully aspirated, and 200 µL of dimethyl sulfoxide (DMSO) was added to each well to solubilize the formazan, after which absorbance was measured at 570 nm using a Bio–Rad microplate reader. Dose-response curves were generated, and IC_50_ values (half–maximal inhibitory concentrations) for each cell line were calculated using GraphPad Prism software (v9.4.1, San Diego, CA, USA). All experiments were conducted in triplicate.

#### Assessment of Synergistic Effects of LEV, RIL, and TMZ

To assess potential synergistic interactions among LEV, RIL, and TMZ, cells were treated with four distinct drug combinations, each formulated based on the individual IC50 values of the respective drugs. CI values were calculated using CompuSyn software (v1.0, ComboSyn Inc., Paramus, NJ, USA), which generates simulations of CI across a range of effect levels (Fa values) and produces Fa-CI plots and isobolograms. Interactions were classified as synergistic (CI < 1), additive (CI = 1), or antagonistic (CI > 1).

#### Cell apoptosis assay

Cell apoptosis was evaluated using an Annexin V-fluorescein isothiocyanate (FITC)/PI staining assay. A total of 5 × 10⁵ cells per well were seeded in six-well plates in RPMI-1640 supplemented with 10% FBS and treated for 24 hours with LEV, RIL, or a TMZ-LEV-RIL combination at concentrations corresponding to the IC50 values determined by the MTT assay. Following treatment, cells were washed with PBS to remove residual medium and dead cells, then harvested by trypsinization using 0.25% trypsin-EDTA. Cells were subsequently washed with PBS (300 × g, 5 min) and resuspended in 500 µL of binding buffer. For staining, 2 µL of Annexin V-FITC and 2 µL of PI were added to the cell suspension, gently mixed, and incubated at room temperature in the dark for 15 minutes. Apoptotic and necrotic cell populations were quantified by flow cytometry (Sysmex-Cyflow), and data were analyzed using FlowJo software (v10.4.0). Apoptotic populations were classified as early apoptosis (Annexin V⁺/PI⁻), late apoptosis (Annexin V⁺/PI⁺), necrosis (Annexin V⁻/PI⁺), and viable cells (Annexin V⁻/PI⁻).

#### RNA extraction, cDNA synthesis, and quantitative real-time PCR

Total RNA was extracted from cultured cells using TRIzol Reagent (Anazol, Iran) according to the manufacturer’s instructions. RNA concentration, purity, and quality were assessed using a NanoDrop 2000 spectrophotometer (Thermo Scientific, USA). Complementary DNA (cDNA) was synthesized from isolated RNA using the PrimeScript RT reagent (YTA, Iran) following the manufacturer’s protocol. Quantitative real-time PCR (RT-qPCR) was performed using AMPLIQON 2× qPCR Master Mix Green (No ROX) on a StepOnePlus Real-Time PCR System (Applied Biosystems, USA). Gene-specific primer sequences used for amplification are provided in Additional file 1: Table S1. Relative gene expression levels were calculated using the 2⁻^ΔΔCt^ method, with *GAPDH* used as the internal reference gene [72].

## Discussion

As the incidence of glial malignancies rises with aging, aging also leads to immunosenescence, which extensively affects the adaptive immune system, particularly T-cell function and metabolism. These factors significantly impact the efficacy of immunotherapy [73, 74]. Ladomersky et al. [75] demonstrated that advanced age increases immunosuppressive factors like IDO1 and PD-L1 in the brain, coinciding with decreased cytolytic T cells. Wu et al. [76] specifically showed that senior patients experience dramatic increases in monocyte-derived macrophages, which inhibit T-cell function and stimulate tumor cell proliferation. In this study, we integrated three classes of EA clocks and stratified GBM cases by EA to conduct an interdisciplinary analysis examining how EA reshapes neuronal activity and remodels the TME of GBM, bridging GBM neuroscience and tumor immunology.

Gliomas comprise a diverse array of cellular components within the TME, with subgroups delineated by distinct cellular states [77]. Epigenomic profiling and deconvolution approaches have proven instrumental in characterizing these subclasses [78], and recent work has underscored the critical role of epigenetic regulation across multiple cancer types [79]. Our deconvolution analysis revealed that EAA GBM harbor a distinct TME characterized by increased malignant content and reduced immune infiltration. In contrast, EAD tumors were associated with worse prognosis and displayed multiple transcriptional modules expressed predominantly by neurons, glial and NPC-like progenitor cells, and immune cells, particularly TAMs. This finding aligns with recent evidence that an epigenetically defined high-neural GBM signature, often enriched in GSCs, predicts adverse outcomes [37].

Using multimodal spatial analysis, we found that glutamatergic signaling emerged as a key communication axis between the neuronal-development niche and the TME. Drexler et al. [37] demonstrated that GBMs with a high neural signature exhibit elevated expression of synaptic integration genes, especially within GSCs of neuronal origin. This heightened neural state promoted neuron-to-glioma synapse formation, suggesting that glioma cells epigenetically recapitulate neural lineage programs to achieve circuit integration. These findings underscore synaptic integration as a key driver of GBM progression and suggest that therapies targeting neuron-glioma networks may offer significant therapeutic potential [80]. A recent study by Nejo et al. [6] demonstrated that inhibiting glutamatergic signaling with Perampanel reversed immunosuppression and enhanced therapeutic efficacy in GBM mouse models. Our results demonstrate that the anti-glutamatergic agents LEV and RIL modulate the expression of key genetic signatures associated with stemness, neuronal activity, and immunosuppression. This molecular reprogramming was accompanied by increased sensitivity to TMZ, highlighting a novel combinatorial strategy to potentially overcome chemoresistance via targeting tumor-neuron crosstalk. According to previous studies, TMZ can induce therapy-driven lineage plasticity, whereby non-stem tumor cells enter a reversible senescent state and subsequently re-enter the cell cycle as more stem-like, invasive, and therapy-resistant progeny [81]. In addition, TMZ generates a chronic stress or hypoxia-like TME that activates stemness-associated programs, including HIF signaling, HMGB1 release, and EMT/mesenchymal-like transitions, thereby contributing to tumor recurrence with increased stemness and intratumoral heterogeneity [82]. A recent study demonstrated that patient-derived GSCs exposed to high-dose TMZ activate specific neuroactive, synaptic-like expression programs [83]. LEV reduces neuronal activity and decreases production of miR-184-3p-enriched exosomes that promote mesenchymal transition of GSCs, thereby increasing radiosensitivity [84]. LEV also enhances TMZ effects by suppressing MGMT expression through histone deacetylase modulation, particularly in GSCs from enhanced lesions [85]. Similarly, RIL synergistically enhances TMZ antitumor effects in MGMT-positive GBM cell lines by suppressing MGMT expression and preventing TMZ-induced MGMT upregulation [86]. Beyond its interaction with TMZ, RIL inhibits glutamate release and mGluR1-dependent signaling, leading to reduced proliferation and viability of both GBM cells and GSCs [87]. In GBM mouse intracranial xenografts, troriluzole monotherapy improved survival, and its combination with an anti-PD-1 antibody demonstrated enhanced efficacy [88]. RIL has been shown to reduce PD-L1 in colorectal models via glutamate inhibition and cGAS/STING activation thereby transiently elevating immune checkpoint expression before enhancing anti-PD-1 responses [89]. Our findings indicate that combined treatment with TMZ and LEV, or TMZ and RIL, counteracts TMZ monotherapy-induced upregulation of stemness and immune evasion markers, supporting the hypothesis that anti-epileptic drugs may enhance the efficacy of TMZ and immunotherapy by reprogramming the GBM tumor microenvironment toward a less aggressive and more differentiated state. We also demonstrated that LEV and RIL enhance the apoptotic effect of TMZ in GBM cells. LEV potentiates the antitumor activity of TMZ in GBM stem cells by inhibiting proliferation and promoting apoptosis via activation of the apoptotic pathway [85]. While RIL induces cell death, including through apoptotic mechanisms, in various cancer types including GBM [90].

We also found that EAD-associated modules, mapped to neurons, was enriched in non-responder patients to PD-1 inhibitor immunotherapy. These modules were also enriched in HFC cases. Our findings align with a recent study by Krishna et al. [42], which demonstrated poorer survival in GBM cases with HFC, as these tumors can more effectively integrate into and exploit neural networks. Similarly, Nejo et al. [6] reported that GBM regions with increased neuronal connectivity exhibit regional immunosuppression and enrichment of anti-inflammatory TAMs. This also aligns with the findings of McFaline-Figueroa et al. [40] who reported that neoadjuvant treatment suppressed cancer proliferation genes and enhanced T-cell/interferon-related gene expression. Our longitudinal analysis further revealed that EAA modules were elevated in initial tumors, whereas EAD signatures were higher in recurrent cases. Prior longitudinal transcriptome and proteome studies have highlighted the dynamic evolutionary patterns in GBM, including a proneural-to-mesenchymal transition, enhanced neuronal activity and synaptogenic pathways in patient-derived xenografts [91], and upregulation of extracellular matrix genes upon recurrence [92]. A recent study by Spitzer et al. [93] utilizing longitudinally matched single-nucleus transcriptomic profiling of GBM samples collected before and after treatment, showed that overall cell-type composition remained largely consistent between primary and recurrent tumors, with a shift toward the MES phenotype observed only in a subset of patients. According to this study, recurrent tumors display decreased malignant cell fractions and a corresponding increase in non-malignant glial and neuronal compartments, a pattern that aligns with our observation of higher neuronal and GSC modules in recurrent GLASS cases.

We acknowledge certain limitations in our study. Utilizing more complex GBM models, such as patient-derived xenografts or specialized systems that better recapitulate tumor-neuron interactions, would enhance the physiological relevance of these findings. Additionally, future research should incorporate Western blotting to evaluate protein levels of marker genes and validate mRNA expression patterns. Finally, the in vitro findings were obtained using established GBM cell lines and require validation in patient-derived models and in vivo systems.

## Conclusion

In conclusion, integrative analysis of EA biomarkers provides a powerful framework for decoding the aging biology of GBM. Our findings identify glutamatergic signaling as a candidate central dysregulated pathway in patients exhibiting EAD, positioning it as a clinically actionable target. Repurposing anti-glutamate agents in combination with TMZ offers promise for synergistic cytotoxic efficacy while mitigating adverse TMZ-associated phenotypes including enhanced stemness, neuronal hyperactivity, and immunosuppression. These insights establish a translational foundation for epigenetically informed, mechanism-driven therapeutic strategies in GBM.

## Availability of data and materials

The TCGA-GBM DNAm, bulk RNA-seq datasets are available at https://portal.gdc.cancer.gov/projects/TCGA-GBM. The IDAT files from Drexler et al. [37], used for external validation, are available in the Gene Expression Omnibus under accession number <u>GSE240704</u>. The bulk RNA-seq datasets supporting the results of this study are publicly available. The GLASS bulk RNA-seq dataset is available from the synapse portal (https://www.synapse.org/#!Synapse:syn17038081/wiki/585622), and the dataset from Zhao et al. [39] is available from NCBI BioProject under accession number PRJNA482620 (https://www.ncbi.nlm.nih.gov/bioproject/PRJNA482620/). Additionally, the IVY-GAP RNA-seq data, is accessible at https://glioblastoma.alleninstitute.org/. The GBMap scRNA-seq dataset is accessible via <u>CELLxGENE</u>. The spatial transcriptomic and metabolomic datasets from the study by Ravi et al. [47] were downloaded from the Dryad Digital Repository (https://doi.org/10.5061/dryad.h70rxwdmj).

## Supplementary Information

**Figure S1.** Analysis of Drexler et al. DNAm as external validation dataset.

**Table S1.** List of primer sequences used in the study.

**Table S2.** List of DMPs between EAD and EAA groups in TCGA-GBM cohort.

**Table S3.** List of DMPs between EAD and EAA groups in Drexler et al. cohort.

## Ethics approval

The Ethics Committee approved this study of Ardabil University of Medical Sciences, Ardabil, Iran (IR.ARUMS.REC.1404.223).

## Clinical trial number

Not applicable.

## Author statement

The manuscript has been read and approved by all the authors, and the requirements for authorship as stated in author structure have been met, and each author believes that the manuscript represents honest work.

## Contribution Details

E.S. and M.M. conceived and planned the study; M.M., M.E., H.A., P.Z., N.S., and M.T. carried out the experiments and collected the available literature; M.M. and E.S. prepared the manuscript. E.S. supervised the study. The manuscript has been read and approved by all named authors.

## Acknowledgments

The authors gratefully acknowledge Khorshid Pharmaceutical Co. (Tehran, Iran) for gifting the active metabolite of RIL.

## Abbreviation

BDNF: Brain-derived neurotrophic factor
CA: chronological age
CT: cellular tumor
DMP: differentially methylated probe
EA: Epigenetic age
EAA: Epigenetic age accelerated
EAD: Epigenetic age decelerated
GBM: Glioblastoma
HBV: hyperplastic blood vessel
HFC: high functional connectivity
IT: Infiltrating tumor
LE: leading edge
LEV: Levetiracetam
MVP: microvascular proliferation
NLGN3: Neuroligin-3
NPC: neural progenitor cell
OPC: oligodendrocyte progenitor cell
PAN: pseudopalisading cells around necrosis
RIL: Riluzole
ST: Spatial transcriptomics
TCGA: The Cancer Genome Atlas
THBS1: thrombospondin-1
TMZ: Temozolomide

---

**Figure. S1.**
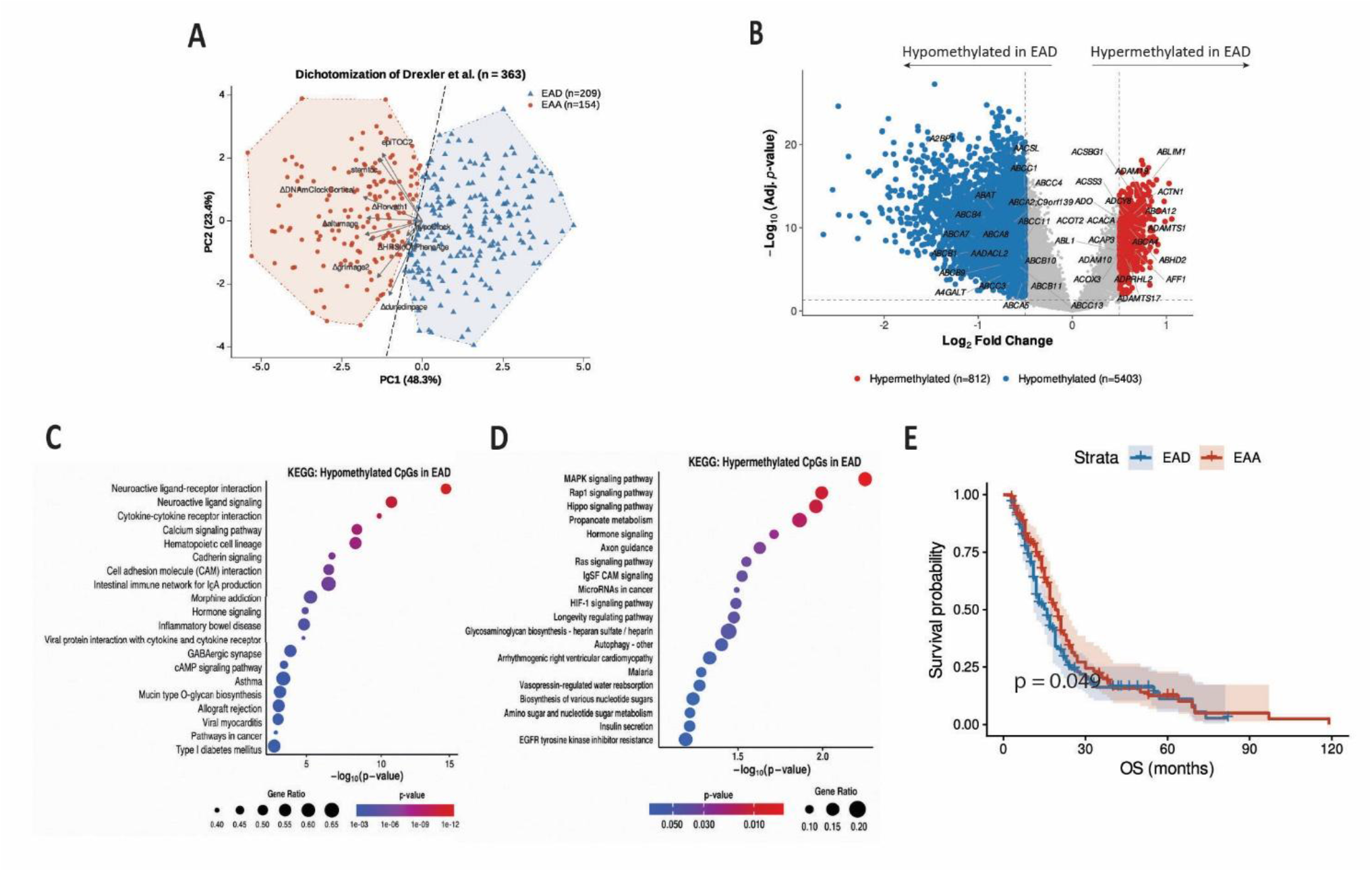
(**A**) PCA-LDA hybrid plot illustrating the separation of EAA and EAD sample groups in Drexler dataset. (**B**) Volcano plot demonstrating differentially methylated genes in EAD vs. EAA. Each dot represents a CpG probe, mapped to its corresponding gene. (**C)** KEGG overrepresentation enrichment analysis of hypormethylated probes in EAD tumors. (**D**) KEGG overrepresentation analysis of hypervvvmethylated probes in EAD tumors. (**E**) Kaplan-Meier curve showing OS of EAA, and EAD cases.

## References

1. Ostrom, Q.T., et al., CBTRUS Statistical Report: Primary Brain and Other Central Nervous System Tumors Diagnosed in the United States in 2016-2020. Neuro Oncol, 2023. 25(12 Suppl 2): p. iv1–iv99.

2. Venkataramani, V., et al., Glutamatergic synaptic input to glioma cells drives brain tumour progression. Nature, 2019. 573(7775): p. 532–538.

3. Venkatesh, H.S., et al., Electrical and synaptic integration of glioma into neural circuits. Nature, 2019. 573(7775): p. 539–545.

4. Venkatesh, H.S., et al., Targeting neuronal activity-regulated neuroligin-3 dependency in high-grade glioma. Nature, 2017. 549(7673): p. 533–537.

5. Taylor, K.R., et al., Glioma synapses recruit mechanisms of adaptive plasticity. Nature, 2023. 623(7986): p. 366–374.

6. Nejo, T., et al., Glioma-neuronal circuit remodeling induces regional immunosuppression. Nature Communications, 2025. 16(1): p. 4770.

7. Thakkar, J.P., et al., Epidemiologic and molecular prognostic review of glioblastoma. Cancer Epidemiol Biomarkers Prev, 2014. 23(10): p. 1985–96.

8. Ostrom, Q.T., et al., CBTRUS Statistical Report: Primary Brain and Other Central Nervous System Tumors Diagnosed in the United States in 2012-2016. Neuro Oncol, 2019. 21(Suppl 5): p. v1–v100.

9. Roa, W., et al., Abbreviated course of radiation therapy in older patients with glioblastoma multiforme: a prospective randomized clinical trial. J Clin Oncol, 2004. 22(9): p. 1583–8.

10. Weller, M., et al., EANO guidelines on the diagnosis and treatment of diffuse gliomas of adulthood. Nat Rev Clin Oncol, 2021. 18(3): p. 170–186.

11. Fane, M. and A.T. Weeraratna, How the ageing microenvironment influences tumour progression. Nature Reviews Cancer, 2020. 20(2): p. 89–106.

12. Stoll, E.A., P.J. Horner, and R.C. Rostomily, The impact of age on oncogenic potential: tumor-initiating cells and the brain microenvironment. Aging Cell, 2013. 12(5): p. 733–41.

13. Ladomersky, E., et al., Advanced Age Increases Immunosuppression in the Brain and Decreases Immunotherapeutic Efficacy in Subjects with Glioblastoma. Clin Cancer Res, 2020. 26(19): p. 5232–5245.

14. Hope, H.C., et al., Age-associated nicotinamide adenine dinucleotide decline drives CAR-T cell failure. Nature Cancer, 2025. 6(9): p. 1524–1536.

15. Segovia, G., A. Porras, A. Del Arco, and F. Mora, Glutamatergic neurotransmission in aging: a critical perspective. Mech Ageing Dev, 2001. 122(1): p. 1–29.

16. Dickstein, D.L., et al., Changes in the structural complexity of the aged brain. Aging Cell, 2007. 6(3): p. 275–84.

17. Horvath, S., DNA methylation age of human tissues and cell types. Genome Biol, 2013. 14(10): p. R115.

18. Ryan, C.P., “Epigenetic clocks”: Theory and applications in human biology. Am J Hum Biol, 2021. 33(3): p. e23488.

19. Yu, M., W.D. Hazelton, G.E. Luebeck, and W.M. Grady, Epigenetic Aging: More Than Just a Clock When It Comes to Cancer. Cancer Res, 2020. 80(3): p. 367–374.

20. Zheng, C., N.A. Berger, L. Li, and R. Xu, Epigenetic age acceleration and clinical outcomes in gliomas. PLoS One, 2020. 15(7): p. e0236045.

21. Mittal, B.B. and D.E. Vaughan, Inclusion of epigenetic age acceleration in oncological trials. Lancet Healthy Longev, 2023. 4(5): p. e185–e186.

22. Delgado-Morales, R., R.C. Agis-Balboa, M. Esteller, and M. Berdasco, Epigenetic mechanisms during ageing and neurogenesis as novel therapeutic avenues in human brain disorders. Clin Epigenetics, 2017. 9: p. 67.

23. Teschendorff, A.E. and S. Horvath, Epigenetic ageing clocks: statistical methods and emerging computational challenges. Nat Rev Genet, 2025. 26(5): p. 350–368.

24. Shireby, G.L., et al., Recalibrating the epigenetic clock: implications for assessing biological age in the human cortex. Brain, 2020. 143(12): p. 3763–3775.

25. Levine, M.E., et al., An epigenetic biomarker of aging for lifespan and healthspan. Aging (Albany NY), 2018. 10(4): p. 573–591.

26. Lu, A.T., et al., DNA methylation GrimAge version 2. Aging (Albany NY), 2022. 14(23): p. 9484–9549.

27. Belsky, D.W., et al., Quantification of the pace of biological aging in humans through a blood test, the DunedinPoAm DNA methylation algorithm. Elife, 2020. 9.

28. Teschendorff, A.E., A comparison of epigenetic mitotic-like clocks for cancer risk prediction. Genome Med, 2020. 12(1): p. 56.

29. Zhu, T., et al., An improved epigenetic counter to track mitotic age in normal and precancerous tissues. Nat Commun, 2024. 15(1): p. 4211.

30. Malta, T.M., et al., Machine Learning Identifies Stemness Features Associated with Oncogenic Dedifferentiation. Cell, 2018. 173(2): p. 338–354 e15.

31. Tepeoglu, M., P. Borcek, O. Ozen, and N. Altinors, Microsatellite Instability in Glioblastoma: Is It Really Relevant in Tumor Prognosis? Turk Neurosurg, 2019. 29(5): p. 778–784.

32. Wang, Q., et al., Tumor Evolution of Glioma-Intrinsic Gene Expression Subtypes Associates with Immunological Changes in the Microenvironment. Cancer Cell, 2017. 32(1): p. 42–56 e6.

33. Verhaak, R.G., et al., Integrated genomic analysis identifies clinically relevant subtypes of glioblastoma characterized by abnormalities in PDGFRA, IDH1, EGFR, and NF1. Cancer Cell, 2010. 17(1): p. 98–110.

34. Ceccarelli, M., et al., Molecular Profiling Reveals Biologically Discrete Subsets and Pathways of Progression in Diffuse Glioma. Cell, 2016. 164(3): p. 550–63.

35. Sidaway, P., CNS cancer: Glioblastoma subtypes revisited. Nat Rev Clin Oncol, 2017. 14(10): p. 587.

36. Torre, M., P.Y. Wen, and J.B. Iorgulescu, The predictive value of partial MGMT promoter methylation for IDH-wild-type glioblastoma patients. Neurooncol Pract, 2023. 10(2): p. 126–131.

37. Drexler, R., et al., A prognostic neural epigenetic signature in high-grade glioma. Nat Med, 2024. 30(6): p. 1622–1635.

38. Pike, S.C., et al., Glioma immune microenvironment composition calculator (GIMiCC): a method of estimating the proportions of eighteen cell types from DNA methylation microarray data. Acta Neuropathol Commun, 2024. 12(1): p. 170.

39. Zhao, J., et al., Immune and genomic correlates of response to anti-PD-1 immunotherapy in glioblastoma. Nat Med, 2019. 25(3): p. 462–469.

40. McFaline-Figueroa, J.R., et al., Neoadjuvant anti-PD1 immunotherapy for surgically accessible recurrent glioblastoma: clinical and molecular outcomes of a stage 2 single-arm expansion cohort. Nat Commun, 2024. 15(1): p. 10757.

41. Barthel, F.P., et al., Longitudinal molecular trajectories of diffuse glioma in adults. Nature, 2019. 576(7785): p. 112–120.

42. Krishna, S., et al., Glioblastoma remodelling of human neural circuits decreases survival. Nature, 2023. 617(7961): p. 599–607.

43. Ruiz-Moreno, C., et al., Harmonized single-cell landscape, intercellular crosstalk and tumor architecture of glioblastoma. bioRxiv, 2022: p. 2022.08.27.505439.

44. Sun, D., et al., Identifying phenotype-associated subpopulations by integrating bulk and single-cell sequencing data. Nat Biotechnol, 2022. 40(4): p. 527–538.

45. Puchalski, R.B., et al., An anatomic transcriptional atlas of human glioblastoma. Science, 2018. 360(6389): p. 660–663.

46. Harwood, D.S.L., et al., Glioblastoma cells increase expression of notch signaling and synaptic genes within infiltrated brain tissue. Nat Commun, 2024. 15(1): p. 7857.

47. Ravi, V.M., et al., Spatially resolved multi-omics deciphers bidirectional tumor-host interdependence in glioblastoma. Cancer Cell, 2022. 40(6): p. 639–655 e13.

48. Jin, S., M.V. Plikus, and Q. Nie, CellChat for systematic analysis of cell-cell communication from single-cell transcriptomics. Nat Protoc, 2025. 20(1): p. 180–219.

49. Moriarty, C., N. Gupta, and D. Bhattacharya, Role of Glutamate Excitotoxicity in Glioblastoma Growth and Its Implications in Treatment. Cell Biol Int, 2025. 49(5): p. 421–434.

50. Pei, Z., et al., Pathway analysis of glutamate-mediated, calcium-related signaling in glioma progression. Biochem Pharmacol, 2020. 176: p. 113814.

51. Oh, M.C., et al., Overexpression of calcium-permeable glutamate receptors in glioblastoma derived brain tumor initiating cells. PLoS One, 2012. 7(10): p. e47846.

52. Wirsching, H.G., et al., Negative allosteric modulators of metabotropic glutamate receptor 3 target the stem-like phenotype of glioblastoma. Mol Ther Oncolytics, 2021. 20: p. 166–174.

53. Kanemaru, K., et al., Spatially resolved multiomics of human cardiac niches. Nature, 2023. 619(7971): p. 801–810.

54. Lee, C.Y., C.C. Chen, and H.H. Liou, Levetiracetam inhibits glutamate transmission through presynaptic P/Q-type calcium channels on the granule cells of the dentate gyrus. Br J Pharmacol, 2009. 158(7): p. 1753–62.

55. Jabbarli, R., et al., How about Levetiracetam in Glioblastoma? An Institutional Experience and Meta-Analysis. Cancers (Basel), 2021. 13(15).

56. Guo, X., et al., Hypoxia-Induced Neuronal Activity in Glioma Patients Polarizes Microglia by Potentiating RNA m6A Demethylation. Clin Cancer Res, 2024. 30(6): p. 1160–1174.

57. Sun, M., et al., The efficacy of temozolomide combined with levetiracetam for glioblastoma (GBM) after surgery: a study protocol for a double-blinded and randomized controlled trial. Trials, 2022. 23(1): p. 234.

58. Doble, A., The pharmacology and mechanism of action of riluzole. Neurology, 1996. 47(6 Suppl 4): p. S233–41.

59. Deheeger, M., M.S. Lesniak, and A.U. Ahmed, Cellular plasticity regulated cancer stem cell niche: a possible new mechanism of chemoresistance. Cancer Cell Microenviron, 2014. 1(5).

60. Beier, D., et al., Temozolomide preferentially depletes cancer stem cells in glioblastoma. Cancer Res, 2008. 68(14): p. 5706–15.

61. Venkatesh, H.S., et al., Neuronal Activity Promotes Glioma Growth through Neuroligin-3 Secretion. Cell, 2015. 161(4): p. 803–16.

62. Pinheiro, K.V., et al., Expression and pharmacological inhibition of TrkB and EGFR in glioblastoma. Mol Biol Rep, 2020. 47(9): p. 6817–6828.

63. Wang, S., et al., Temozolomide promotes immune escape of GBM cells via upregulating PD-L1. Am J Cancer Res, 2019. 9(6): p. 1161–1171.

64. Thrush, K.L., Higgins-Chen, A. T., Liu, Z. & Levine, M. E R methylCIPHER: A Methylation Clock Investigational Package for Hypothesis-Driven Evaluation & Research. 2022. DOI: 10.1101/2022.07.13.499978.

65. de Lima Camillo, L.P., *pyaging: a Python-based compendium of GPU-optimized aging clocks*. Bioinformatics, 2024. 40(4).

66. Patro, R., et al., Salmon provides fast and bias-aware quantification of transcript expression. Nat Methods, 2017. 14(4): p. 417–419.

67. Morabito, S., et al., hdWGCNA identifies co-expression networks in high-dimensional transcriptomics data. Cell Rep Methods, 2023. 3(6): p. 100498.

68. Cerami, E., et al., The cBio cancer genomics portal: an open platform for exploring multidimensional cancer genomics data. Cancer Discov, 2012. 2(5): p. 401–4.

69. Kleshchevnikov, V., et al., Cell2location maps fine-grained cell types in spatial transcriptomics. Nat Biotechnol, 2022. 40(5): p. 661–671.

70. Causer, A., et al., SpaMTP: Integrative Statistical Analysis and Visualisation of Spatial Metabolomics and Transcriptomics data. bioRxiv, 2024: p. 2024.10.31.621429.

71. Phipson, B., J. Maksimovic, and A. Oshlack, missMethyl: an R package for analyzing data from Illumina’s HumanMethylation450 platform. Bioinformatics, 2016. 32(2): p. 286–8.

72. Livak, K.J. and T.D. Schmittgen, Analysis of relative gene expression data using real-time quantitative PCR and the 2(-Delta Delta C(T)) Method. Methods, 2001. 25(4): p. 402–8.

73. Han, S., et al., Age-associated remodeling of T cell immunity and metabolism. Cell Metab, 2023. 35(1): p. 36–55.

74. Kaiser, M., et al., Immune Aging and Immunotherapy in Cancer. Int J Mol Sci, 2021. 22(13).

75. Ladomersky, E., et al., The Coincidence Between Increasing Age, Immunosuppression, and the Incidence of Patients With Glioblastoma. Front Pharmacol, 2019. 10: p. 200.

76. Wu, S., et al., Integrative analysis of single-cell transcriptomics reveals age-associated immune landscape of glioblastoma. Front Immunol, 2023. 14: p. 1028775.

77. Neftel, C., et al., An Integrative Model of Cellular States, Plasticity, and Genetics for Glioblastoma. Cell, 2019. 178(4): p. 835–849 e21.

78. Singh, O., D. Pratt, and K. Aldape, Immune cell deconvolution of bulk DNA methylation data reveals an association with methylation class, key somatic alterations, and cell state in glial/glioneuronal tumors. Acta Neuropathol Commun, 2021. 9(1): p. 148.

79. Terekhanova, N.V., et al., Epigenetic regulation during cancer transitions across 11 tumour types. Nature, 2023. 623(7986): p. 432–441.

80. Winkler, F., et al., Cancer neuroscience: State of the field, emerging directions. Cell, 2023. 186(8): p. 1689–1707.

81. Wang, P., et al., Temozolomide promotes glioblastoma stemness expression through senescence-associated reprogramming via HIF1alpha/HIF2alpha regulation. Cell Death Dis, 2025. 16(1): p. 317.

82. Gao, X.Y., et al., Temozolomide Treatment Induces HMGB1 to Promote the Formation of Glioma Stem Cells via the TLR2/NEAT1/Wnt Pathway in Glioblastoma. Front Cell Dev Biol, 2021. 9: p. 620883.

83. Marei, H.E., et al., Distinct molecular pathways leading to dosage-dependent temozolomide resistance in GBM stem cells. Cancer Cell International, 2026. 26(1): p. 118.

84. Guo, X., et al., Neuronal Activity Promotes Glioma Progression by Inducing Proneural-to-Mesenchymal Transition in Glioma Stem Cells. Cancer Res, 2024. 84(3): p. 372–387.

85. Scicchitano, B.M., et al., Levetiracetam enhances the temozolomide effect on glioblastoma stem cell proliferation and apoptosis. Cancer Cell Int, 2018. 18: p. 136.

86. Yamada, T., et al., Riluzole enhances the antitumor effects of temozolomide via suppression of MGMT expression in glioblastoma. J Neurosurg, 2021. 134(3): p. 701–710.

87. Sperling, S., et al., Riluzole: a potential therapeutic intervention in human brain tumor stem-like cells. Oncotarget, 2017. 8(57): p. 96697–96709.

88. Medikonda, R., et al., Synergy between glutamate modulation and anti-programmed cell death protein 1 immunotherapy for glioblastoma. J Neurosurg, 2022. 136(2): p. 379–388.

89. Liang, B., et al., Riluzole Enhancing Anti-PD-1 Efficacy by Activating cGAS/STING Signaling in Colorectal Cancer. Mol Cancer Ther, 2025. 24(1): p. 131–140.

90. Blyufer, A., et al., Riluzole: A neuroprotective drug with potential as a novel anti–cancer agent (Review). Int J Oncol, 2021. 59(5).

91. Kim, K.H., et al., Integrated proteogenomic characterization of glioblastoma evolution. Cancer Cell, 2024. 42(3): p. 358–377 e8.

92. Hoogstrate, Y., et al., Transcriptome analysis reveals tumor microenvironment changes in glioblastoma. Cancer Cell, 2023. 41(4): p. 678–692 e7.

93. Spitzer, A., et al., Deciphering the longitudinal trajectories of glioblastoma ecosystems by integrative single-cell genomics. Nat Genet, 2025. 57(5): p. 1168–1178.

