## Supplementary material for "Glioblastoma Tumors with Decelerated Epigenetic Aging Are Characterized by Glutamatergic Neuronal Activity and Stemness": Table S1

| **Gene** | **Forward/Reverse** | **Sequence** |
| --- | --- | --- |
| *SOX2* | Forward | GCTACAGCATGATGCAGGACCA |
|  | Reverse | TCTGCGAGCTGGTCATGGAGTT |
| *EZH2* | Forward | TTGTTGGCGGAAGCGTGTAAAATC |
|  | Reverse | TCCCTAGTCCCGCGCAATGAGC |
| *TUBB3* | Forward | TCAGCGTCTACTACAACGAGGC |
|  | Reverse | GCCTGAAGAGATGTCCAAAGGC |
| *GALC* | Forward | ACTCTCACCACTGGTCGCAAAG |
|  | Reverse | GATCAGCAAAGTTTGGAGCTTCAC |
| *NTRK2 (TrKB)* | Forward | ACAGTCAGCTCAAGCCAGACAC |
|  | Reverse | GTCCTGCTCAGGACAGAGGTTA |
| *NLGN3* | Forward | GTCTGGTTCACTGCCAACTTGG |
|  | Reverse | CCGTCATTATCCGCTAAGTCCTC |
| *THBS1* | Forward | GCTGGAAATGTGGTGCTTGTCC |
|  | Reverse | CTCCATTGTGGTTGAAGCAGGC |
| *SYP* | Forward | TCGGCTTTGTGAAGGTGCTGCA |
|  | Reverse | TCACTCTCGGTCTTGTTGGCAC |
| *CD274 (PD-L1)* | Forward | ATCAAGTCCTGAGTGGTAAG |
|  | Reverse | GAGGTAGTTCTGGGATGA |
